# Higher-order metabolic interactions among oral commensals gate pathobiont expansion through succinate-driven pH modulation

**DOI:** 10.64898/2026.11.04.764248

**Authors:** Dam Soh, Aadil H. Bhat, Lu Li, Matthew Smardz, Carol Parker, Erin R. Tiede, Brynn Nelson, Valerie A. Frerichs, Troy D. Wood, Laertis Ikonomou, Hung Ton-That, Patricia I. Diaz

## Abstract

Despite advances in defining microbiome shifts associated with inflammatory mucosal diseases, the ecological interactions that constrain or facilitate pathobiont expansion remain poorly understood. Using a six-species synthetic oral microbial community and factorial interaction screening, we identified a higher-order interaction governing the growth of the periodontal pathobiont *Porphyromonas gingivalis*. The core community member *Veillonella parvula* was the strongest positive determinant of *P. gingivalis* growth, but this effect was substantially reduced by the health-associated commensal *Actinomyces oris*. *A. oris*-mediated antagonism resulted from extracellular acidification and was enhanced under anaerobic conditions. Genetic dissection of *A. oris* fermentation revealed that lactate dehydrogenase (*ldh1*) contributes to anaerobic lactate production, but *ldh1* deletion only partially relieved *P. gingivalis* inhibition. Instead, succinate, produced through a succinate dehydrogenase/fumarate reductase pathway (*sdh2*), emerged as the principal driver of acidification and *P. gingivalis* suppression. At high cell densities, however, *V. parvula* consumed both lactate and succinate, reversing acidification and restoring conditions permissive for *P. gingivalis* growth. To assess the relevance of these interactions in vivo, we analyzed subgingival microbiome from 1,355 adults, including 555 with detectable *P. gingivalis*. Among these individuals, *P. gingivalis* abundance was negatively associated with *Actinomyces* abundance after adjustment for periodontitis severity, whereas this association was attenuated at high *V. parvula* abundance, consistent with the interaction observed in vitro. Together, these findings identify succinate as a previously unrecognized mediator of interbacterial antagonism and reveal how higher-order metabolic interactions among oral commensals regulate pathobiont expansion through pH modulation, with protective effects shaped by oxygen availability and metabolic cross-feeding.

## Introduction

Understanding the determinants that govern the colonization and expansion of pathobionts within microbial communities is critical for defining strategies to maintain microbiome health and prevent disease. The colonization and growth of potentially pathogenic microbial species within a community depend on interactions with resident microorganisms and with the local physicochemical and host environments. Microorganisms can facilitate the growth of other species through cooperative resource utilization^1,2^ and metabolic cross-feeding^3,4^, while resource competition^5^ and antimicrobial production^6,7^ represent exclusion mechanisms. Microbes also influence growth of others by modifying their shared environment. Changes in pH, oxygen availability, or other physicochemical properties can create conditions that favor some organisms while restricting those with different physiological requirements^8,9^. Disentangling these effects in surveys of natural microbiomes remains challenging because observed patterns of microbial abundance reflect the combined outcome of all these factors acting simultaneously.

The oral microbiome represents an accessible, well-characterized ecosystem suitable for interrogating the drivers of pathobiont expansion. Dental plaque communities derive nutrients from host fluids, including saliva and gingival exudates, and encounter different oxygen gradients depending on their location relative to the gingival margin and their stage of biofilm development. These environmental gradients alter microbial metabolism and may therefore change the nature and consequences of interactions among community members. Understanding these interactions is particularly relevant in the context of the oral disease periodontitis, a chronic inflammatory condition characterized by destruction of the tooth-supporting tissues and associated with a dysbiotic subgingival microbiome^10–12^. *Porphyromonas gingivalis* (Pg) is a Gram-negative, anaerobic, asaccharolytic pathobiont strongly associated with periodontitis and implicated in disease development and progression^13^. However, Pg colonization appears highly variable as it is not detected in all individuals with severe periodontitis, its abundance varies considerably among those colonized, and it can also be detected at low abundance in healthy individuals^14^. We recently identified cell-density-dependent growth as one factor constraining Pg colonization within dental biofilms^14^. This restriction can be overcome through an interaction with *Veillonella parvula* (Vp), a core species abundant both in homeostasis and disease. In pairwise cultures, Vp provides a soluble factor that enables Pg growth from low cell density inocula and aids its oral colonization in a mouse model^14^. Although these and other pairwise studies have uncovered interactions that facilitate Pg establishment in dental plaque^15,16^, whether health-associated microbiome members can actively restrain Pg expansion and how the broader microbial and environmental context influence these effects remains largely unexplored.

Synthetic microbial communities provide an opportunity for systematic identification of inter-species interactions that influence pathobiont growth and dissecting the mechanisms underlying them^17,18^. Previously, using a 6-species synthetic oral microbial community comprising Pg and prevalent plaque commensals — *Actinomyces oris*, *Streptococcus sanguinis*, *Fusobacterium nucleatum*, and *Rothia dentocariosa*, in addition to Vp — we showed that the presence of Vp is required for the optimal growth of Pg in this community, validating the observations from pairwise cultures^14^. However, the behavior of individual species within multispecies communities can also be shaped by higher-order interactions involving three or more species, which can modify the effects of individual community members and generate outcomes that cannot be predicted from pairwise relationships alone^18,19^. Thus, although Vp is necessary to support Pg growth within the six-species community, the reductionist approaches used previously could not resolve additional interactions among community members that may modulate Pg growth.

Here, we use this six-species synthetic community together with a Design of Experiments (DOE) factorial methodology to systematically evaluate higher-order interspecies interactions that regulate Pg growth within a community context. This approach uncovered a conditional antagonistic interaction in which the health-associated commensal *Actinomyces oris* (Ao) restricts Pg growth through anoxia-dependent extracellular acidification. Molecular dissection of the Ao fermentation products responsible for pH modulation and Pg inhibition, identified succinate production as a critical driver of extracellular acidification and Pg inhibition, whereas Vp counteracted this effect at high cell densities by consuming Ao-produced organic acids thereby restoring a pH permissive to Pg growth. We then tested whether the ecological relationships predicted by these experimentally defined interactions were evident in the human subgingival microbiome. Across a population-scale cohort, *Actinomyces* abundance was negatively associated with that of Pg, with this relationship attenuated at high Vp abundance. Together, these findings demonstrate that health-associated taxa can actively create conditions that restrict pathobiont growth and reveal how this antagonism is shaped by environmental conditions and metabolic interactions among community members.

## Results

### Factorial screening reveals interspecies interactions that shape Pg growth

To evaluate interspecies interactions that influence *Porphyromonas gingivalis* (Pg) growth within a defined polymicrobial context, we utilized a previously established six-species oral community model comprising Pg and five early-colonizing commensals: *Actinomyces oris* (Ao)*, Veillonella parvula* (Vp)*, Fusobacterium nucleatum* (Fn), *Streptococcus sanguinis* (Ss), and *Rothia dentocariosa* (Rd)^14^. These commensals were selected based on their high prevalence and abundance in dental plaque and their ability to colonize prior to Pg establishment^20,21^. This community, grown in mucin serum, has been shown to support Pg growth from a low cell-density inoculum, whereas Pg fails to grow in monoculture under similar inoculation conditions^14^. During community growth, the early-colonizing commensals increase in biomass within the first 48 hours, followed by Pg expansion, which reaches late exponential phase by day 4^14^. Previous pair-wise experiments further identified Vp as the only individual member of this community capable of supporting low cell density Pg growth^14^.

To determine whether interactions involving several community members influenced Pg growth, we employed a half-fraction factorial experiment design comprising 17 conditions, to systematically assess the effects of the five commensals, individually and in combinations, on Pg growth rate (h ¹) during 4 days of culture (Figure 1a). Under all conditions, the commensals grew effectively achieving their maximum biomass after 24 hours (Supplemental Figure S1a). Factorial analysis identified Vp as the strongest positive determinant of Pg growth (Figure 1b), with its effect exceeding the Bonferroni-adjusted significance threshold (Supplementary Figure S1b), consistent with our previous findings from pairwise cultures^14^. In contrast, Ao exerted a significant negative effect on Pg growth. Importantly, the growth-promoting effect of Vp was strongly dependent on the presence of Ao, resulting in a significant negative Ao × Vp interaction (Figure 1c). Together, these findings suggest that Pg growth is governed not only by the presence of growth-supporting species such as Vp, but also by interspecies interactions that determine whether their growth-promoting effects are realized.

**Figure 1.**
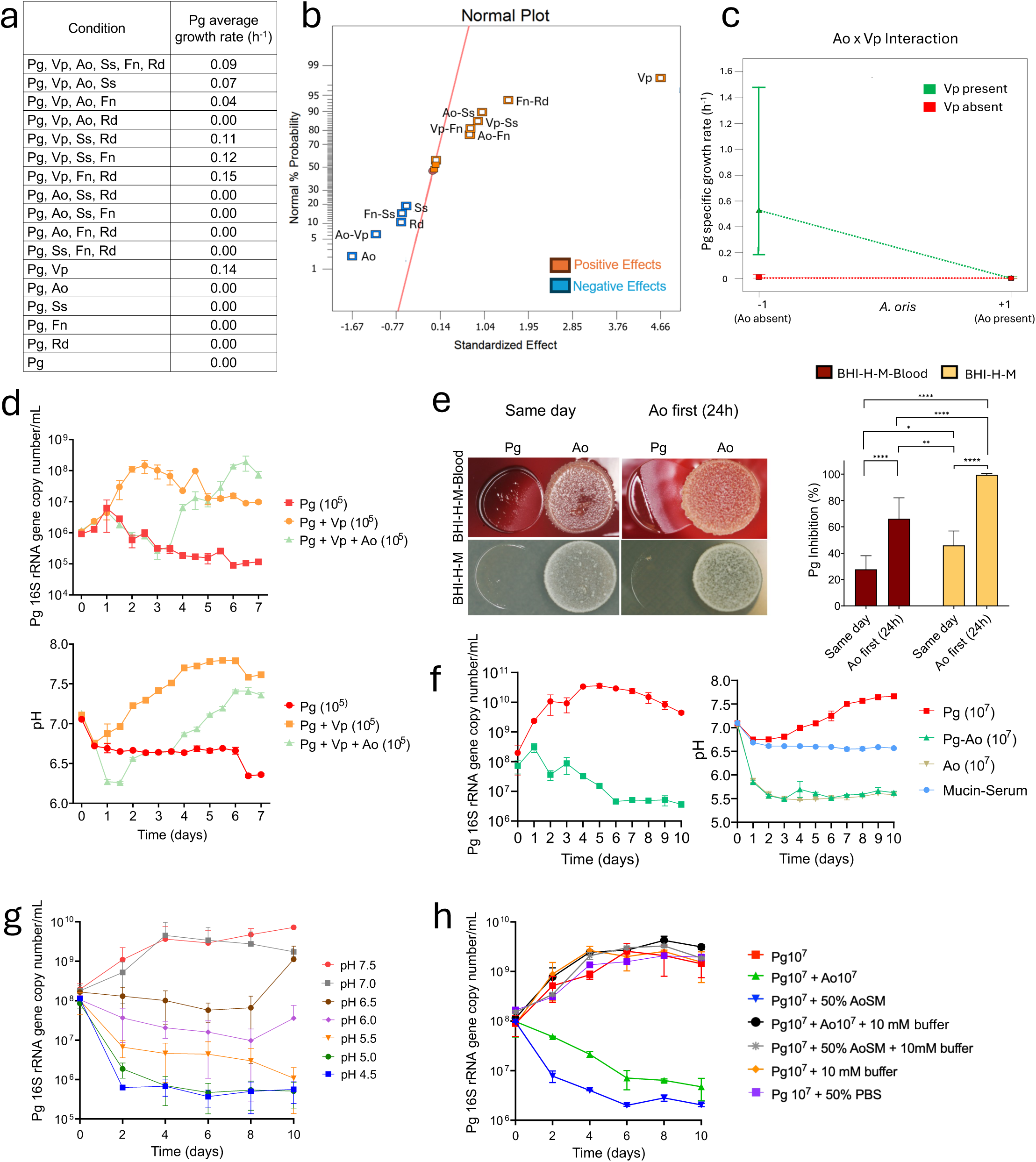
Interspecies interactions and pH-mediated inhibition of *Porphyromonas gingivalis* (Pg) by *Actinomyces oris* (Ao) in a polymicrobial community. (a) Half factorial screening of the effects of five species and their interactions on Pg growth rate in mucin-serum. The 16 conditions were evaluated in duplicate and in two independent experiments. The five species included *Actinomyces oris* (Ao) *Veillonella parvula* (Vp), *Streptococcus sanguinis* (Ss), *Fusobacterium nucleatum* (Fn) and *Rothia dentocariosa* (Rd). Pg growth rate was calculated from logarithmic phase in cultures evaluated for 4 days. (b) Normal probability plot of standardized effects of individual species and their interactions on Pg growth rate. Orange and blue squares indicate positive and negative effects, respectively. The red line provides a reference for effects consistent with random variation. (c) Model-predicted effect of the two-factor interaction between Ao and Vp on Pg growth rate (h ¹). Red and green symbols indicate Vp absence and presence, respectively. Dashed lines show model predictions. Error bars represent standard deviation or 95% confidence interval. (d) Growth of Pg inoculated at low density (10 cells/mL) alone, with Vp, or with Vp and Ao in mucin–serum medium. The upper and lower panels show Pg biomass measured by qPCR and the corresponding culture pH, respectively. (e) Adjacent competitive growth of Ao and Pg on two different agar media. Pg was spotted simultaneously with Ao or after Ao had grown anaerobically for 24 h. Differences in Pg growth inhibition were assessed using one-way ANOVA followed by Fisher’s least significant difference (LSD) test, and multiple comparisons correction via the Benjamini–Hochberg method. * *P* < 0.05, *** *P* < 0.001, **** *P* < 0.0001. (f) Growth of Pg inoculated at high density (10 cells/mL) alone or with Ao in mucin–serum medium. The left and right panels show Pg biomass measured by qPCR and the corresponding culture pH, respectively. Ao monocultures and the medium-only control provided pH references. The same Pg inoculum density and medium were used in g and h. (g) Growth of Pg in mucin-serum adjusted to different initial pH values. (h) Growth of Pg with Ao or 50% (v/v) Ao spent medium under unbuffered or buffered conditions. Fresh and spent media were buffered with 10 mM potassium phosphate (pH 7.3).

### Ao inhibits growth of Pg through environmental acidification

To validate the interactions identified by the factorial analysis, we conducted co-culture experiments with Ao, Vp, and Pg and monitored Pg biomass over 7 days. Consistent with our previous findings, Pg inoculated at low cell density failed to grow, whereas the addition of Vp supported robust Pg growth (Figure 1d). Inclusion of Ao with Vp and Pg, suppressed Pg growth through day 4, coinciding with a decline in the culture pH (Figure 1d). However, this suppression was transient as Pg growth recovered after day 4, concomitant with an increase in pH. The suppression of Pg growth was not due to inhibition of Vp by Ao, as Vp biomass was comparable between Pg-Vp and Pg-Vp-Ao co-cultures (Supplementary figure S1c).

To determine whether Ao directly inhibits Pg under conditions where Pg can grow independently, we first performed agar-based competition spot assays. Ao and Pg were either inoculated simultaneously onto supplemented BHI agar with or without blood, or Ao was allowed to grow for 24 h before Pg inoculation. Ao inhibited Pg growth in both agar media, with significantly greater inhibition when Ao was spotted 24 hours before Pg inoculation, suggesting a priority effect, and when blood was absent (Figure 1e). We next examined the Pg-Ao interaction in liquid mucin-serum cultures using a high-density Pg inoculum, which supports autonomous growth. Under these conditions, Ao again inhibited Pg growth with this inhibition coinciding with a rapid decline in culture pH to values close to 5.5 and with Pg unable to recover (Figure 1f).

Given previous reports of Pg sensitivity to acidic environments^22^, we assessed growth of Pg in mucin-serum medium adjusted to different initial pH levels ranging from 4.5 to 7.5. As seen in Figure 1g, growth of Pg was pH-dependent with robust growth at pH ≥7.0, and progressively impaired growth under more acidic conditions. We next tested whether the inhibitory effect of Ao on Pg growth resulted from acidification of the extracellular environment by supplementing the medium with 10 mM potassium phosphate buffer (pH 7.3). High-density Pg grew robustly in monoculture but showed marked inhibition when co-cultured with either Ao or Ao cell-free spent media. Buffering the medium completely rescued Pg growth under both conditions, altogether demonstrating that Ao inhibits Pg by lowering the extracellular pH beyond levels permissive for robust Pg growth (Figure 1h).

### Oxygen availability modulates Ao-mediated acidification and inhibition of Pg

Ao is a facultative anaerobe commonly found in both supragingival and subgingival plaque^23,24^, where differences in oxygen availability require metabolic flexibility. Previous models of Ao metabolism suggest that under oxygen-limited conditions, pyruvate flux from the Embden– Meyerhof–Parnas (EMP) pathway is shunted toward lactate (pKa of ∼3.86) and succinate (diprotic, pKa ∼4.21 and pKa ∼5.64) production to maintain redox homeostasis via NADH reoxidation^25,26^. In contrast, aerobic metabolism is predicted to favor production of the weaker acid acetate (pKa ∼4.76) (Supplementary Figure S2). Therefore, we hypothesized that Ao-mediated acidification and the resulting inhibition of Pg growth would be oxygen-dependent, with greater acidification and Pg inhibition occurring under anaerobic conditions.

To test this hypothesis, we first assessed acidification by Ao colonies grown under different redox conditions on phenol red agar. Under aerobiosis, Ao colonies produced minimal color change on the agar, whereas anaerobic growth resulted in a pronounced yellow transition indicative of acidification to pH <6.0 (Figure 2a). We next investigated whether this oxygen-dependent acidification influences the antagonistic activity of Ao against Pg. In competitive spot assays, Ao pre-cultured anaerobically for 24 hours exerted significantly greater inhibition against Pg compared to aerobically-grown Ao (Figure 2b). Consistently, in liquid mucin-serum monocultures, Ao dropped the pH to 5.5 during anaerobic growth, while aerobically the pH remained close to neutral values (7.0). Since lactate is a predicted end-product of anaerobic fermentation, we next quantified lactate in liquid mucin-serum monocultures of Ao, which showed lactate was detected exclusively under anaerobic conditions (Figure 2c).

**Figure 2.**
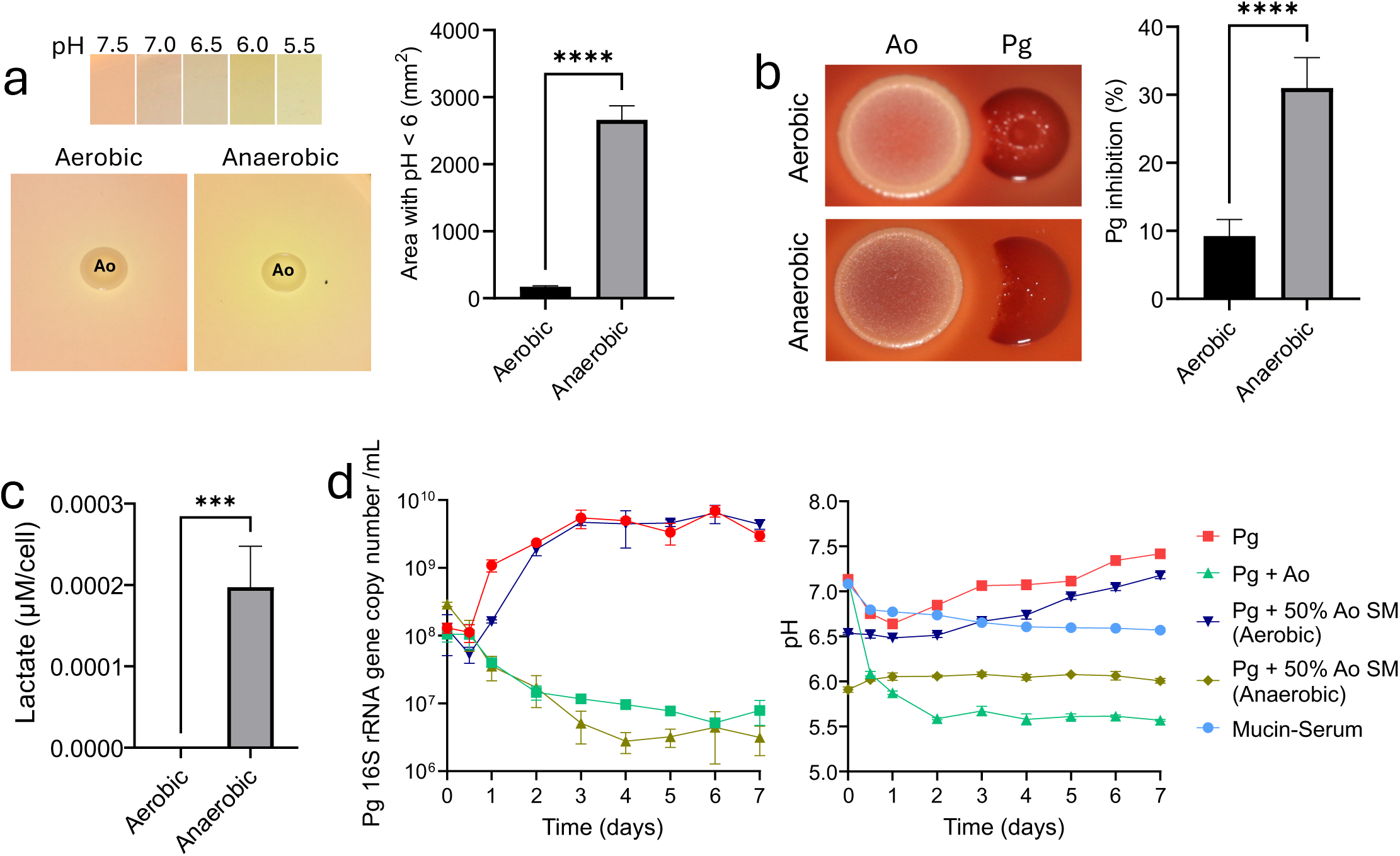
Oxygen availability modulates Ao metabolism, medium acidification, and subsequent inhibition of Pg. (a) Acidification of the surrounding medium by Ao grown on BHI agar containing phenol red for 24 h under aerobic or anaerobic conditions. The pH color scale is shown above the images. The bar plot quantifies the area surrounding Ao colonies with pH < 6.0. (b) Pg growth inhibition in spot competition assays on BHI blood agar supplemented with hemin and menadione. Ao was pre-cultured under aerobic or anaerobic conditions for 24 h followed by Pg spotting and anaerobic incubation. The bar plot quantifies Pg growth inhibition. (c) Quantification of lactate in Ao spent medium after 24 h of aerobic or anaerobic growth. Lactate concentrations were normalized to Ao viable biomass as determined by colony-forming units (CFU). (d) Growth of Pg inoculated at high density (10 cells/mL) in mucin–serum medium supplemented with 50% (v/v) Ao spent medium (SM) collected from aerobic or anaerobic cultures in late exponential phase. Controls included Pg–Ao co-cultures and Pg monocultures. The left and right panels show Pg abundance and the corresponding culture pH, respectively. The medium-only control provides a pH reference. Pg abundance was quantified by qPCR and expressed as 16S rRNA gene copies/mL. Differences between aerobic and anaerobic conditions in a–c were assessed using unpaired Student’s t-tests. *** *P* < 0.001, **** *P* < 0.0001.

Next, spent medium collected from Ao cultures grown under aerobic or anaerobic conditions was tested for its ability to inhibit Pg. Cultures containing spent medium derived from anaerobically-grown Ao maintained an acidic pH (∼6.0) and strongly inhibited Pg growth, whereas cultures with spent medium from aerobically grown Ao had a pH slightly above 6.5 and allowed Pg proliferation (Figure 2d). These findings demonstrate that Ao-mediated acidification and inhibition of Pg are strongly dependent on oxygen availability.

### Genetic dissection of Ao fermentation reveals ldh1-dependent lactate production contributes to, but does not fully account for, Ao antagonism of Pg

Although early physiological models conceptualized the fermentation pathways of Ao^26^, the specific genetic determinants governing these metabolic activities remain undefined. Therefore, we next aimed to molecularly identify the genomic regions responsible for acid production in Ao and establish the genes and products responsible for its antagonism toward Pg. We identified two paralog genes in the Ao genome annotated as L-lactate dehydrogenase (Ldh), with 64.8% nucleotide sequence identity and 17.9% protein sequence identity, predicted to encode for enzymes with different functions (Figure 3a). One gene, hereafter referred to as *ldh1*, was predicted to encode a canonical fermentative NAD-dependent Ldh (EC 1.1.1.27), which converts pyruvate to lactate while oxidizing NADH to NAD^+^ (EC 1.1.1.27)^27,28^, while the second gene, hereafter referred to as *ldh2*, was predicted to function as a quinone-dependent Ldh (EC 1.1.5.15), which catalyzes lactate oxidation to pyruvate coupled to quinone reduction to quinol (EC 1.1.5.15)^29^. Thus, the two paralogs are predicted to mediate opposing reactions in lactate metabolism, with Ldh1 favoring lactate production and Ldh2 lactate utilization. Based on these predicted functions, we hypothesized that *ldh1* would mediate anaerobic lactate production and contribute to Ao-dependent inhibition of Pg.

**Figure 3.**
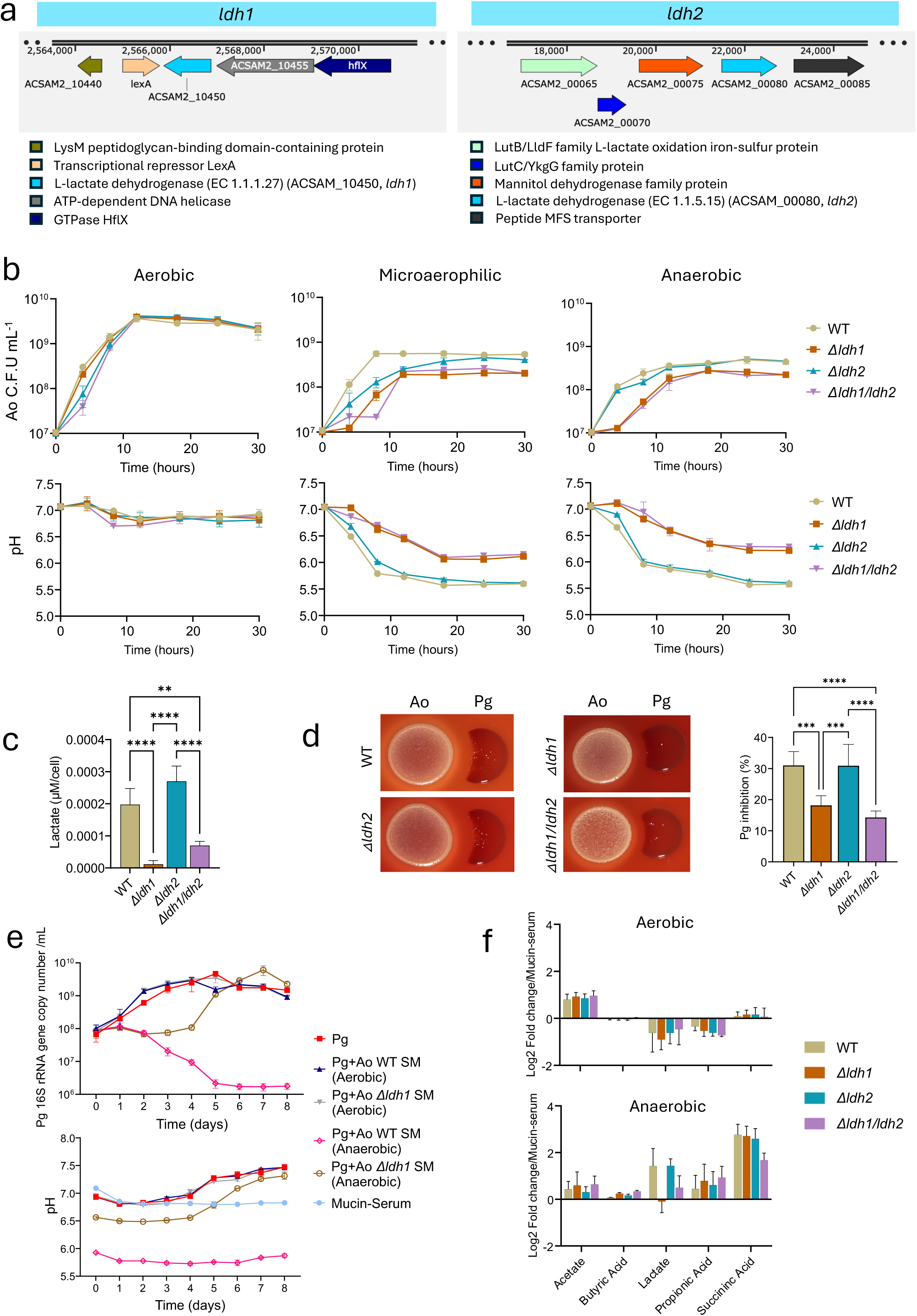
*ldh1* of Ao is essential for anaerobic lactate production and contributes to inhibition of *Pg*. (a) Genomic neighborhoods of Ao *ldh1* (ACSAM2_10450, EC 1.1.1.27, left) and *ldh2* (ACSAM2_00080, EC 1.1.5.15, right). Arrows indicate open reading frames and direction of transcription, with gene annotations and designated locus tags color-coded according to predicted function. Chromosomal coordinates are indicated in base pairs (bp). (b) Growth and medium acidification of Ao wild-type (WT) and lactate dehydrogenase (*ldh*) mutant strains grown in mucin-serum under different redox conditions. The upper panels show Ao biomass over time and lower panels show corresponding culture pH. (c) Extracellular lactate production by Ao WT and *ldh* mutant strains cultured under anaerobic conditions. Total lactate was quantified from cell-free culture supernatants using a colorimetric assay and normalized to CFU counts (μM cell^-1^). (d) Pg growth inhibition in spot competition assays on BHI blood agar. Ao WT and *ldh* mutant strains were spotted 24 h, and allowed to grow anaerobically, before Pg inoculation. (e) Growth of Pg in mucin–serum medium supplemented with spent medium (SM) from Ao WT or Δ*ldh1* cultures grown aerobically or anaerobically. Pg monoculture was included as a control. The upper and lower panels show Pg abundance and the corresponding culture pH, respectively. The medium-only control provides a pH reference. Pg abundance was quantified by qPCR and expressed as 16S rRNA gene copies/mL. (f) Targeted LC-MS analysis of organic end-products produced by Ao WT and mutant strains under aerobic and anaerobic growth in mucin-serum. Values represent log fold changes in metabolite concentrations relative to uninoculated medium. Error bars in b–f indicate standard deviations. Differences among strains in c and d were assessed using one-way ANOVA followed by Fisher’s least significant difference (LSD) test. We corrected for multiple comparisons by controlling the false discovery rate using the Benjamini–Hochberg method. ** *P* < 0.01, *** *P* < 0.001, **** *P* < 0.0001.

To test this hypothesis, we employed an in-frame deletion strategy to generate single (*Δldh1* or *Δldh2*) and double-deletion mutants (*Δldh1/ldh2*) of Ao. Under aerobic conditions, all strains displayed growth kinetics comparable to the wild-type and did not acidify the culture medium (Figure 3b). In contrast, under anaerobiosis, the Δ*ldh1* and Δ*ldh1/ldh2* mutants exhibited impaired growth and reduced acidification, decreasing the pH to values between 6.0–6.5 compared with a pH of 5.5 for wild-type Ao (Figure 3b). Similar growth and acidification phenotypes were consistent across the two media tested, BHI and mucin-serum (Supplementary Figure S3). Consistent with the anaerobic phenotypes, colorimetric lactate quantification revealed a nearly complete loss of lactate in the Δ*ldh1* and substantial reduction in the Δ*ldh1/ldh2* strain (Figure 3c), confirming a major role for *ldh1* in anaerobic lactate production.

We next determined whether loss of *ldh1* altered Ao-mediated inhibition of Pg. In competitive spot assays, both the Δ*ldh1* and Δ*ldh1/ldh2* mutants exhibited significantly reduced, but not completely abolished, inhibition of Pg compared with wild-type Ao and the Δ*ldh2* mutant (Figure 3d). Furthermore, spent media from aerobically-grown Ao wild-type or Δ*ldh1* failed to inhibit Pg proliferation in liquid culture. In contrast, spent medium from anaerobically-grown Δ*ldh1* suppressed Pg growth for 4 days, after which growth resumed, representing a partial loss of the sustained inhibitory activity observed with wild-type Ao (Figure 3e). Collectively, these findings establish *ldh1* as the major determinant of anaerobic lactate production in Ao and demonstrate that *ldh1*-dependent metabolism contributes to extracellular acidification and Pg inhibition. However, the effect of *ldh1* deletion on Pg inhibition was partial, indicating that additional products contribute to the antagonistic activity of Ao.

To identify additional metabolic end-products that could account for the residual acidification observed anaerobically in the *ldh1* mutant, we performed targeted LC–MS metabolomic profiling of Ao wild-type, Δ*ldh1*, Δ*ldh2*, and Δ*ldh1/ldh2* cultures grown under aerobic and anaerobic conditions. Under aerobiosis, the metabolite profiles of all strains were characterized predominantly by production of acetate, with a small accumulation of succinate (Figure 3f). Under anaerobiosis we found Ao wild-type produces a broader mixture of organic acids, including acetate, lactate, propionate, and succinate. Among these, succinate showed the greatest accumulation across all strains, remaining abundant in the Δ*ldh1* mutant despite the loss of lactate production.

### Sdh2-dependent succinate production by Ao mediates acidification and inhibition of Pg

Prompted by the marked accumulation of succinate during anaerobic growth, we interrogated the Ao genome to identify the genetic basis of succinate production. We identified two distinct loci annotated as succinate dehydrogenase/fumarate reductase complexes (EC 1.3.5.1). Both gene clusters encode a flavoprotein catalytic subunit, an iron–sulfur subunit, and a membrane-anchoring cytochrome *b* subunit, and are located at distinct, unlinked chromosomal regions (Figure 4a). Pairwise sequence alignment of the catalytic flavoprotein subunits (AXE84_07215 and AXE84_01890) revealed 66.7% nucleotide sequence identity and 55.3% protein sequence identity, indicating that they represent two divergent homologs. To determine their functional roles, we generated single deletion mutants targeting the catalytic subunit of each complex, designated *Δsdh1* (ACSAM2_04915) and *Δsdh2* (ACSAM2_12050), as well as a double deletion mutant *Δsdh1/sdh2*.

**Figure 4.**
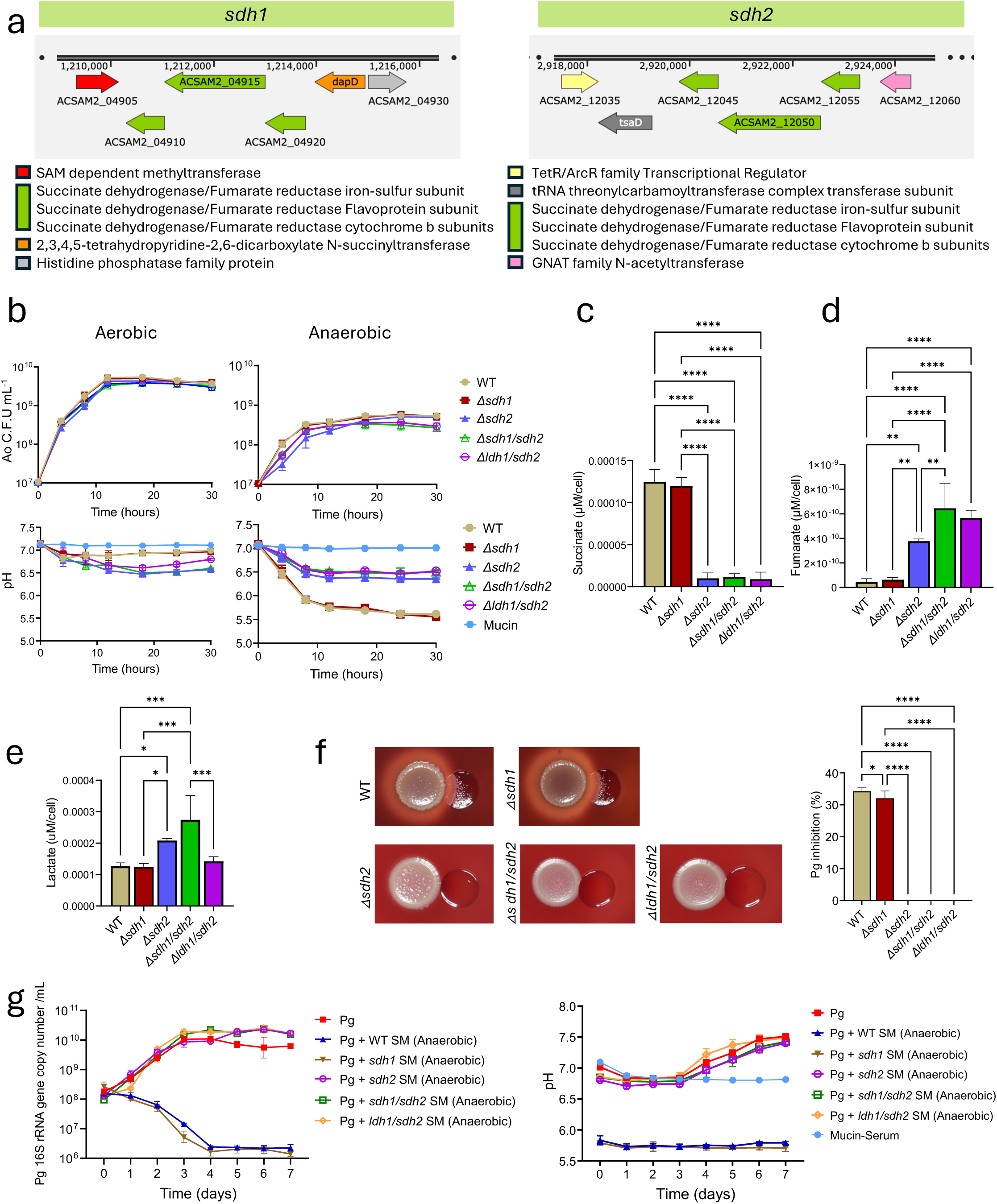
Ao *sdh2* is essential for anaerobic succinate production and inhibition of Pg. (a) Genomic neighborhoods of Ao *sdh1* and *sdh2*, encoding paralogous succinate dehydrogenase/fumarate reductase heterotrimeric complexes (SDH/FRD; EC 1.3.5.1): iron– sulfur subunit (ACSAM2_04910 and ACSAM2_12045), flavoprotein subunit (ACSAM2_04915 and ACSAM2_12050), and cytochrome *b* subunit (ACSAM2_04920 and ACSAM2_12055). Arrows indicate open reading frames and direction of transcription, color-coded by predicted functional category. Chromosomal coordinates are indicated in base pairs (bp). (b) Growth and medium acidification of Ao WT and succinate dehydrogenase (*sdhA*) mutant strains grown in mucin-serum under different redox conditions. The upper panels show Ao growth; the lower panels show corresponding culture pH. (c-e) Extracellular succinate (c), fumarate (d) and lactate (e) in cell-free culture supernatants of Ao WT and *sdh* mutant strains grown anaerobically. Metabolite concentrations were measured by enzymatic colorimetric assays and normalized to total cell counts. (f) Pg growth inhibition in adjacent spot competition assays on BHI blood agar. Ao WT and *sdh* mutant strains were spotted and grown anaerobically for 24 h before Pg inoculation. (g) Growth of Pg in cell-free spent medium from anaerobic cultures of Ao WT or *sdh* mutant strains. Pg monoculture in mucin–serum medium served as a control. The left and right panels show Pg abundance and the corresponding culture pH, respectively. The medium-only control provides a pH reference. Pg abundance was quantified by qPCR and expressed as 16S rRNA gene copies/mL. Error bars in b–g indicate standard deviations. Differences among strains in c–f were assessed using one-way ANOVA followed by Fisher’s least significant difference (LSD) test. We corrected for multiple comparisons by controlling the false discovery rate using the Benjamini–Hochberg method. * *P* < 0.05, ** *P* < 0.01, *** *P* < 0.001, **** *P* < 0.0001.

Growth of all *sdh* mutants was comparable to wild-type Ao under both aerobic and anaerobic conditions (Figure 4b). However, the Δ*sdh2* and Δ*sdh1/sdh2* mutants showed markedly reduced anaerobic acidification, reaching a final pH of approximately 6.5 compared with ∼5.5 for wild-type Ao, whereas Δ*sdh1* retained an acidification phenotype similar to the wild-type. Consistent with these phenotypes, anaerobic succinate accumulation was markedly reduced in the Δ*sdh2* and Δ*sdh1/sdh2* mutants and was accompanied by reciprocal accumulation of fumarate (Figure 4c-d). Aerobically, the Δ*sdh2* and Δ*sdh1/sdh2* mutants showed the opposite phenotype, acidifying the medium slightly more than wild-type (pH ∼6.5 versus ∼7.0) and accumulating small amounts of succinate (Supplementary Figure S4b). This reversal is consistent with the bidirectional nature of succinate dehydrogenase/fumarate reductase complexes in which Sdh2 reduces fumarate to succinate anaerobically, so its loss abolishes succinate production, whereas under aerobiosis it oxidizes succinate to fumarate, and its loss permits succinate to accumulate. Together, these findings identify *sdh2*, but not *sdh1*, as the principal locus responsible for fumarate reduction to succinate during anaerobic growth and demonstrate that *sdh2*-dependent succinate production is a major contributor to Ao-mediated environmental acidification.

Disruption of the succinate-producing pathway also altered lactate metabolism. Loss of *sdh2* was associated with increased anaerobic lactate production, with a statistically significant increase observed in the Δ*sdh2* and Δ*sdh1/sdh2* mutants, which produced significantly more lactate than wild-type Ao (Figure 4e). Deletion of *sdh1* alone had no effect on lactate accumulation, suggesting that impairment of Sdh2-dependent fumarate reduction redirects anaerobic metabolism toward lactate production.

To further examine the relative contributions of lactate and succinate to anaerobic acidification, we generated Δ*ldh1/sdh1* and Δ*ldh1/sdh2* double mutants. Although deletion of *sdh1* alone did not impair anaerobic growth, the Δ*ldh1/sdh1* double mutant failed to grow under anaerobic conditions and was therefore excluded from subsequent metabolic analyses. In contrast, under anaerobic conditions, the Δ*ldh1/sdh2* mutant exhibited robust growth but attenuated acidification similar to that of the Δ*sdh2* single mutant (Figure 4b), together with a comparable reduction in succinate accumulation and reciprocal accumulation of fumarate (Figure 4c-d). Surprisingly, despite deletion of *ldh1*, the Δ*ldh1/sdh2* mutant accumulated lactate to levels comparable to wild-type Ao (Figure 4e), suggesting compensatory mechanisms for anaerobic lactate production. Importantly, lactate accumulation in the Δ*ldh1/sdh2* mutant did not restore medium acidification, thereby uncoupling lactate production from the low-pH phenotype. Furthermore, all *sdh* single and double mutants showed increased pyruvate relative to wild-type Ao (Supplementary Figure S4a), consistent with broader redistribution of central carbon metabolism following disruption of fumarate reduction. Together, these findings demonstrate that acidification more closely tracks Sdh2-dependent succinate production than lactate accumulation, identifying succinate as the major determinant of the acidic phenotype.

We next determined whether Sdh2-dependent succinate production was required for Ao-mediated inhibition of Pg. In competitive spot assays, all *sdh2*-deletion mutants completely lost their capacity to inhibit Pg growth whereas wild-type Ao and the Δ*sdh1* mutant retained inhibitory activity (Figure 4f, Supplementary Figure 4c). Consistently, spent media harvested from anaerobic cultures of Ao wild-type and *Δsdh1* completely suppressed Pg growth, whereas spent media from anaerobically-grown *sdh2*-deletion mutants failed to inhibit Pg (Figure 4g). Concurrent culture pH monitoring demonstrated that spent media from *sdh2*-deficient mutants remained above 6.5, allowing Pg to proliferate. Also, as expected, none of the spent media collected under aerobic conditions exerted any suppressive effect on Pg (Supplementary Figure S4d). Together, these findings demonstrate that Sdh2-dependent succinate production is the critical pathway mediating the inhibition of Pg.

### Vp alleviates Ao-mediated inhibition of Pg through organic acid cross-feeding

Previous experiments showed that co-culture growth of low-cell density Pg with Ao and Vp is initially suppressed but Pg is eventually able to overcome the inhibition and grow (Figure 1d). Since Vp is known to utilize lactate and succinate as carbon and energy sources^30^, we evaluated whether this commensal could under certain conditions abrogate Ao inhibition of Pg. We therefore tested whether Vp influenced Pg in high-density inoculum co-cultures, where Pg growth is independent of Vp, and the high biomass of Vp may allow it to rapidly rescue Pg from Ao inhibition. This experiment showed that in the absence of Vp, co-culture with either Ao or Ao spent medium resulted in marked and sustained inhibition of Pg growth but addition of Vp completely abrogated the Ao inhibitory effect (Figure 5a).

**Figure 5.**
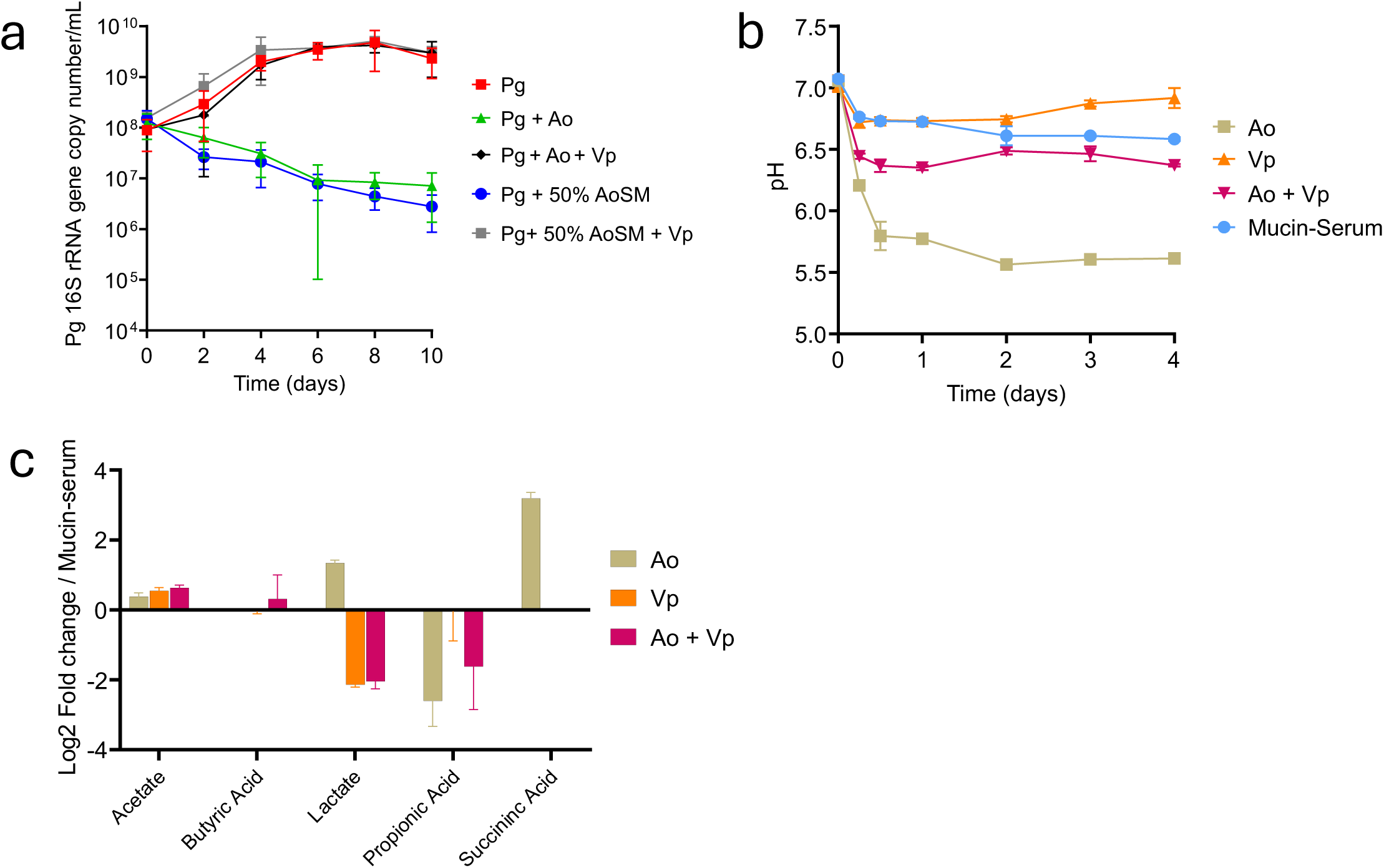
Vp rescues Pg growth through consumption of Ao-derived organic acids. (a) Growth of Pg inoculated at high density (10 cells/mL) in mucin–serum medium with Ao or spent medium harvested from anaerobic Ao cultures, in the presence or absence of Vp. Pg abundance was quantified by qPCR and expressed as 16S rRNA gene copies/mL. (b) Culture pH dynamics of Ao and Vp monocultures and Ao–Vp co-cultures grown anaerobically in mucin–serum medium. Uninoculated medium served as a control. (c) Colorimetric quantification of extracellular lactate in cell-free supernatants from Ao and Vp monocultures and Ao-Vp co-culture after 24 h of anaerobic growth. Differences among cultures were assessed using one-way ANOVA followed by Fisher’s least significant difference (LSD) test. Multiple comparisons correction was performed by controlling the false discovery rate using the Benjamini–Hochberg method. *** *P*< 0.001, **** *P* < 0.0001. (d) Targeted LC-MS metabolomic profiling of organic acids in cell-free supernatants from Ao and Vp monocultures and Ao-Vp co-culture grown anaerobically in mucin-serum medium. Values represent log fold changes in metabolite concentrations relative to uninoculated medium.

We next examined the manner in which Vp modified the medium pH when in coculture with Ao. Addition of Vp restored culture pH toward neutrality in Ao–Vp cocultures (Figure 5b), suggesting that Vp alleviates Ao-induced acid stress. We therefore evaluated the ability of Vp to consume organic acids produced by Ao. LC–MS profiling revealed that Vp completely metabolized Ao-derived lactate and succinate (Figure 5c). Together, these findings indicate that Vp mitigates Ao-mediated Pg inhibition by metabolically removing organic acids that drive acidification, thereby creating conditions permissive for Pg growth.

### A negative association between Actinomyces and Pg in the human subgingival microbiome is attenuated in individuals with high Vp abundance

To investigate whether the interspecies interactions among Pg, Ao and Vp observed in vitro were reflected in human subgingival microbial communities, we analyzed previously generated 16S rRNA gene sequencing data from subgingival plaque of 1,355 individuals^31^, spanning the full range of periodontal health, from no periodontitis to severe periodontitis, as defined by the CDC/AAP epidemiological classification^32^. Subgingival plaque is a primary ecological niche for Pg and provides the low-oxygen conditions relevant to the Ao-mediated inhibition observed experimentally. Ao and Vp were highly prevalent across periodontal disease categories, whereas Pg prevalence was approximately 35% in individuals with no, mild, or moderate periodontitis and increased to 55.3% in those with severe periodontitis (Supplementary Figure S5a). Co-occurrence analysis across all samples showed higher Pg prevalence in Vp-positive than Vp-negative samples. Within both Vp strata, Pg prevalence was lower in Ao-positive samples though neither comparison reached statistical significant by Fisher’s exact test (Supplementary Figure S5c).

Because species detection alone may not capture abundance-dependent interactions, we next examined relative abundance relationships. All three species showed substantial variation in abundance when present in subgingival communities (Figure 6a). Ao abundance was higher in health and decreased with periodontitis severity, whereas Pg showed the opposite trend. However, some individuals with no or mild periodontitis harbored relatively high levels of Pg, suggesting that its expansion is influenced by factors beyond disease status. Based on our experimental findings that Vp counteracts Ao-mediated acidification, we hypothesized that if Pg and Ao showed a negative association even after adjusting for disease severity, this relationship would weaken at higher Vp abundance. We tested this prediction in Pg-positive samples (n=555), where we observed that Ao and Pg abundances were negatively correlated in the low/medium-Vp group (r = −0.258, *P* < 0.001), whereas the correlation was weaker and not statistically significant in the high-Vp group (r = −0.128, *P* = 0.083) (Figure 6b). In regression models using log□□-transformed relative abundances and adjusting for CDC periodontal disease classification, Ao was negatively associated with Pg in the low/medium-Vp reference group (β = −0.131, *P* <0.001). Although the estimated association was weaker at high Vp abundance, the Ao × high-Vp interaction was not statistically significant (β = 0.067, *P* = 0.183) (Figure 6b).

**Figure 6.**
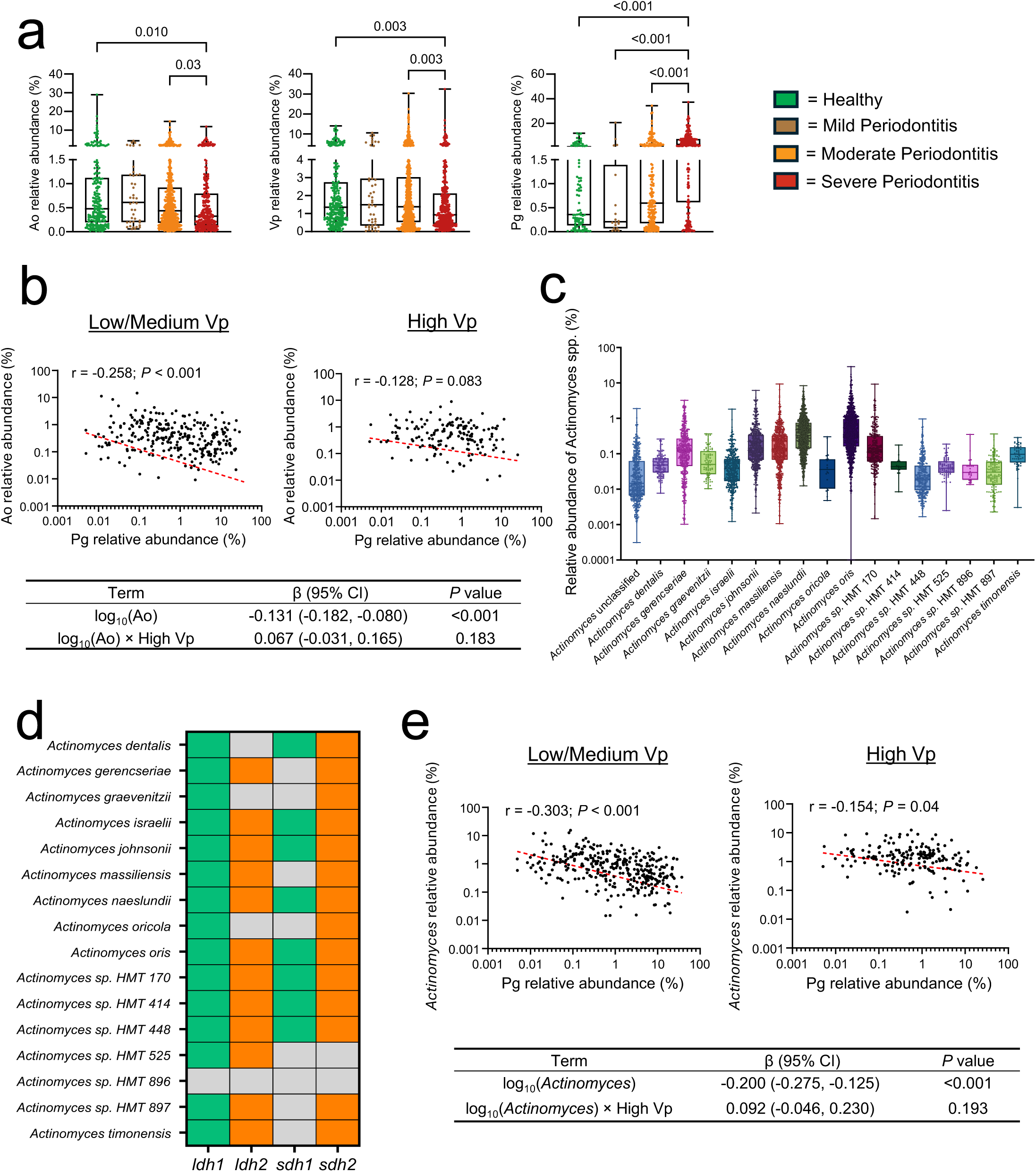
Pg is negatively associated with Ao and the genus *Actinomyces* in the human subgingival microbiome. (a) relative abundance of Ao, Vp, and Pg in subgingival plaque samples from 1,355 participants across periodontal disease severity groups, as determined by 16S rRNA gene sequencing. Differences among clinical groups were assessed using Kruskal–Wallis tests followed by Dunn’s pairwise comparisons, with the Benjamini–Hochberg adjustment to control the false discovery rate. (b) Correlations between Ao and Pg relative abundances in Pg-positive samples (n=555), stratified into low/medium-Vp (left) and high-Vp groups (right). Red dashed lines show linear fits; Pearson’s correlation coefficients (r) and *P* values are indicated. The table below reports selected coefficients (β), 95% confidence intervals and P values for Ao abundance and its interaction with high-Vp, from a CDC-adjusted multiple linear regression model of Pg abundance. The interaction coefficient represents the difference in slope between the two Vp groups. (c) Relative abundances of 16 *Actinomyces* species identified in subgingival plaque samples. (d) Genomic presence of *ldh1, ldh2, sdh1* and *sdh2* across the 16 identified *Actinomyces* species. Colored cells indicate gene presence; grey cells indicate absence. (e) Correlations between genus-level *Actinomyces* and Pg relative abundances, with plot annotations and the corresponding regression table presented as in b.

The pattern observed for Ao prompted examination of whether similar associations extended to the *Actinomyces* genus. Like Ao, total *Actinomyces* relative abundance varied across disease categories and showed a decrease with periodontal disease severity (Supplemental Figure S5b). Taxonomic profiling identified 16 *Actinomyces* species in subgingival plaque samples (Figure 6c). Interrogation of genomes revealed that most of these species encoded *ldh1* and *sdh2*, genes implicated in Pg inhibition in our experimental system (Figure 6d), motivating examination of whether total *Actinomyces* abundance captured a more pronounced negative association with Pg than Ao alone. A consistent pattern was observed when assessing *Actinomyces* abundance in relation to Pg. The relative abundance of *Actinomyces* negatively correlated with Pg in the low/medium Vp group (r = −0.303, *P* < 0.001) with this relationship weakening in the high-Vp stratum (r = −0.154, *P* = 0.040) (Figure 6e). In the corresponding CDC-adjusted regression model, *Actinomyces* abundance was negatively associated with Pg abundance in the low/medium-Vp reference group (β = −0.200, *P* < 0.001). The interaction with high-Vp tends to attenuate this negative effect but remained non-significant (β = 0.092, *P* = 0.193) (Figure 6e). Together, these exploratory population-level patterns are directionally consistent with the experimental model in which Ao/*Actinomyces* inhibits Pg and Vp counteracts this inhibition.

## Discussion

Our findings reveal that the capacity of the resident microbiota to constrain pathobiont expansion is not an intrinsic property of individual commensal species, but emerges from their metabolic activity within a particular environmental and community context. In this study, we combined a factorial synthetic-community screen, targeted genetic dissection, and analysis of a large human cohort to resolve how oral commensals influence the growth of a periodontal pathobiont. We show that the antagonism of the health-associated commensal Ao on the pathobiont Pg is not a fixed, pairwise trait but a conditional, higher-order interaction — reversed in the presence of Vp and dependent on oxygen availability — that traces mechanistically to a dominant pH-modulating metabolite. These findings reframe *Actinomyces*, and potentially other homeostasis-associated commensals, not as passive markers of oral health but as active, context-dependent participants in colonization resistance against pathobionts.

This conditional antagonism illustrates why community-level phenotypes cannot be reliably predicted by the simple summation of pairwise interactions^18,33–35^. To explore species interactions in a polymicrobial community we employed a Design of Experiments (DOE) statistical framework, an approach developed for industrial process optimization but rarely applied to microbial ecology. Fractional factorial designs allow the main effects of multiple species and their interactions to be estimated simultaneously from a limited number of community configurations, making systematic interrogation of higher-order effects tractable without culturing every possible combination. Recent work by Ishizawa and colleagues^18^ has established the value of beyond-pairwise observation for understanding community assembly using a seven-member synthetic community. Quantification of all possible 127 subset combinations and mathematical modeling showed that interaction coefficients derived from three-member combinations predict community structure far better than pairwise coefficients alone. In our study, we used the factorial screen not as an endpoint but as a discovery tool, identifying a significant Ao × Vp interaction term and then dissecting it genetically to the metabolic loci and end-products responsible. By constructing targeted deletions in Ao, we established that its antagonism on Pg is dictated by anaerobic fermentation mainly through *sdh2* and to a lesser extent through *ldh1*. This demonstrates that complex, higher-order microbial interactions discovered via DOE can be resolved into discrete genetic determinants. Moreover, our follow up experiments added several layers of conditionality to the initially discovered interaction, with the redox environment determining whether Ao acts as an antagonist, and with succinate and lactate consumption by Vp abrogating the antagonistic effect of Ao of Pg.

Environmental niche modification represents a mechanism of colonization resistance, in which commensals restrict pathobiont establishment not via contact-dependent antagonism or direct nutrient competition, but by creating an inhospitable microenvironment^8,36,37^. Extracellular pH serves as an ecological filter across diverse human microbiomes^37–39^. In the gut, for example, resident anaerobes exclude invading *Escherichia coli* through organic acid-mediated acidification below pH 5.5^9^. In the oral cavity, environmental acidification has been extensively characterized in supragingival plaque in the context of dental caries^39,40^, but the role of pH modulation in governing subgingival polymicrobial biofilm assembly and pathobiont colonization has remained largely uncharacterized. Here, we confirmed that Pg is highly sensitive to acidic stress, growing optimally under neutral-to-alkaline conditions, which are characteristic of inflamed, diseased periodontal pockets where Pg readily expands^22,41^. The timing of colonization also mattered, with Ao established 24 h before Pg inhibiting it substantially more than Ao introduced simultaneously, indicating a priority effect. Because acidification constrains Pg establishment rather than eliminating an existing population, communities in which Ao is already resident and Vp is scarce would be expected to resist Pg colonization most strongly, whereas Pg arriving into a Vp-rich community, or established before Ao, should be comparatively unaffected. Moreover, the proteolytic virulence of Pg, including gingipain expression and activity require near-neutral pH^42,43^. Since virulence is compromised at pH values above those that restrict growth, even modest commensal-driven acidification would therefore attenuate Pg pathogenicity in communities where the organism persists. pH may therefore act as an ecological constraint on both the growth and virulence of Pg in vivo, with microbial partnerships such as that with Vp enabling Pg to overcome this limitation.

Our findings identify succinate as the main metabolic effector driving this exclusionary acidification. Although lactate is conventionally viewed as the primary driver of oral acidogenesis due to its prominent role in dental caries^44,45^, Ao employs a branched anaerobic fermentation pathway that yields distinct organic acids in the absence of oxygen, including substantial amounts of succinate through fumarate reduction. Intriguingly, despite comparable per-cell secretion of lactate and succinate, genetic dissection resolved succinate as the dominant physiological inhibitor of Pg. This difference may reflect distinct proton balances associated with lactate and succinate fermentation, resulting in different net effects on extracellular pH. Additionally, succinate may be particularly effective at maintaining inhibitory conditions. Unlike lactic acid (pKa 3.86), succinic acid is diprotic, with a second pKa of 5.64, closer to the pH threshold of 6.5 below which Pg growth is restricted. Thus, accumulated succinate may provide greater buffering capacity against alkalinization driven by Pg peptide catabolism, prolonging growth inhibition. Consistent with this interpretation, in an unbuffered medium chemically adjusted to pH 6.5, without ongoing Ao-mediated succinate replenishment, Pg eventually overcame the initial acid stress and grew after a prolonged lag.

The identification of succinate as an ecological barrier also provides a striking mechanistic counterpoint to its function in the human gut. There, succinate is normally held at low concentrations by commensal consumption but following antibiotic treatment it accumulates transiently, and the pathobiont *Clostridioides difficile* exploits this spike to expand, converting succinate to butyrate through an inducible pathway^46^. In the subgingival microbiome, the same intermediate may operate in an opposing ecological capacity. Unconsumed succinate accumulation suppresses the pathobiont Pg through acidification, whereas its utilization by Vp relieves this inhibition. Succinate consumption is therefore protective in the gut, where it denies a usable substrate to the pathobiont, but permissive in the subgingival crevice, where it removes an acid barrier that Pg, being asaccharolytic, can neither exploit nor tolerate. This divergence demonstrates that the ecological role of a shared microbial metabolite is not an intrinsic molecular property, but depends on pathobiont physiology and the microbiome context.

Our genetic dissection of Ao fermentation revealed metabolic plasticity and functional redundancy when central electron sinks are perturbed. Under anaerobic conditions, glycolytic flux depends on continuous NAD regeneration to prevent redox collapse. In wild-type Ao, this redox equilibrium appears to be maintained primarily through two diverging routes, Ldh1-mediated reduction of pyruvate to lactate and Sdh2-dependent reduction of fumarate to succinate. The direction of Sdh2-dependent flux is itself oxygen-dependent, as under anaerobiosis its loss abolished succinate production, whereas under aerobiosis it permitted succinate to accumulate, consistent with the bidirectional nature of succinate dehydrogenase/fumarate reductase complexes. When the dominant succinate branch is disrupted in the Δ*sdh2* and Δ*sdh1/sdh2* strains, cells redirect reducing equivalents and accumulated upstream pyruvate toward the Ldh1 pathway, resulting in lactate overproduction. Resilience of this fermentative network was also evident in the Δ*ldh1/sdh2* double mutant. Despite lacking both its primary lactate dehydrogenase (*ldh1*) and primary fumarate reductase (*sdh2*), this strain grew robustly and restored extracellular lactate secretion to wild-type levels. This phenotype implies a bypass route, possibly mediated by derepressed activity of the secondary lactate dehydrogenase (*ldh2*) operating in reverse under elevated intracellular pyruvate and NADH/NAD pools, or by alternative oxidoreductases capable of substrate-level electron disposal. Because Ldh2 is predicted to be quinone-dependent, such reverse flux would additionally require a sufficiently reduced quinone pool, and direct flux measurements will be needed to establish this.

In contrast to Δ*ldh1/sdh2*, the Δ*ldh1/sdh1* mutant failed to grow under anaerobic conditions, even though Sdh1 contributes negligibly to net succinate accumulation and its single deletion produced no change in succinate, fumarate, lactate, final pH, or Pg inhibition. Metabolite profiling nonetheless revealed that Δ*sdh1* accumulated anaerobic pyruvate to levels exceeding those of Δ*sdh2*, indicating that Sdh1 does carry appreciable flux despite leaving end-product profiles unchanged. We therefore propose that Sdh1 provides a secondary route for electron disposal, dispensable while Ldh1 is intact but essential once the principal NADH sink is removed. Together, these genetic interactions illustrate that Ao does not depend on a rigid linear fermentation route, but rather on a flexible network capable of rerouting carbon and reducing equivalents under redox constraints.

This metabolic rewiring is intimately governed by oxygen tension, rendering the protective capacity of Ao strictly context-dependent. As a facultative anaerobe possessing metabolic plasticity, Ao shifts from oxidative respiration to mixed-acid fermentation under anoxia, markedly enhancing organic acid excretion and localized acidification to exert potent antagonism against Pg. In contrast, under aerobic conditions, respiration attenuates net proton accumulation, substantially abolishing this inhibitory barrier. This oxygen-dependent behavior would confer a different functionality to Ao in the oxygenated supragingival environment versus the more anaerobic subgingival crevice. At a broader level, these dynamics illustrate that taxonomic composition alone cannot predict microbiome function. Ao is not an intrinsically protective commensal, but a conditional environmental regulator whose barrier function emerges only when local physicochemical conditions drive fermentative acidification.

Our findings identify *V. parvula* as an important modulator of the local chemical environment that counteracts *A. oris*-mediated inhibition through metabolic cross-feeding. By consuming lactate and succinate produced by *A. oris*, *V. parvula* alleviates extracellular acidification and restores a pH permissive for *P. gingivalis* growth. However, this effect depends on *V. parvula* cell density and the duration of the interaction. At the low initial cell densities used in our factorial screen, *A. oris* established an inhibitory acidic environment, resulting in a significant negative *A. oris* × *V. parvula* interaction despite the strong growth-promoting effect of *V. parvula* on *P. gingivalis*. In extended three-species cultures, also inoculated at low density, *P. gingivalis* growth recovered after day 4, whereas in follow-up experiments a high initial density of *V. parvula* prevented inhibition from the outset. These findings suggest that *V. parvula* must reach sufficient abundance for organic acid consumption to overcome acid production by *A. oris*, making the outcome of this interaction dependent on initial cell density and time. Interestingly, analysis of human subgingival microbiomes revealed a similar trend, with the negative association between *Actinomyces* and *P. gingivalis* abundance appearing weaker in individuals with high *V. parvula* abundance. Although the statistical interaction was not significant, this pattern is compatible with the density-dependent modulation observed experimentally. Importantly, this mechanism extends the previously described role of *V. parvula* in promoting *P. gingivalis* growth through a diffusible growth-initiating signal¹. Thus, *V. parvula* can facilitate pathobiont expansion through two distinct mechanisms: directly, by providing a growth-promoting signal, and indirectly, by modifying the local environment to relieve the inhibition imposed by another commensal.

Together, our findings illustrate how pathobiont expansion can be governed by higher-order metabolic interactions that dynamically modify the physicochemical environment. Rather than acting as fixed promoters or inhibitors of pathogen growth, resident microbiota can exert context-dependent effects determined by both community composition and environmental conditions.

## Material and methods

### Bacterial strains and culture conditions

*Porphyromonas gingivalis* ATCC 33277 (Pg) was maintained short-term on brain heart infusion (BHI) agar supplemented with 0.4 mg mL^-1^L-cysteine·HCl, 5 µg mL^-1^hemin, 5 µg mL menadione, and 5% (v/v) defibrinated sheep blood. Pg starter cultures were grown in liquid BHI-H-M medium which consisted of 37 g L^-1^ of BHI supplemented with 0.4 mg mL^-1^ L-cysteine·HCl, 5 µg mL^-1^ hemin, and 5 µg mL^-1^ menadione. *Fusobacterium nucleatum* ATCC 10953 (Fn) was maintained on BHI agar containing 0.4 mg mL L-cysteine·HCl and 5% defibrinated sheep blood, with liquid starter cultures prepared in BHI supplemented solely with 0.4 mg mL^-1^L-cysteine·HCl. *Streptococcus sanguinis* SK36 (Ss), *Rothia dentocariosa* ATCC 17931 (Rd), wild-type *Actinomyces oris* MG1 (Ao), and its mutant derivatives (Δ*ldh1,* Δ*ldh2,* Δ*ldh1/*Δ*ldh2,* Δ*sdh1,* Δ*sdh2,* Δ*sdh1/*Δ*sdh2,* Δ*ldh1/*Δ*sdh1* and Δ*ldh1/*Δ*sdh2*) were maintained on BHI agar and cultured in liquid BHI. *Veillonella parvula* PK1910 (Vp) was maintained on BHI agar supplemented with 0.4 mg mL^-1^L-cysteine·HCl, 14.9 µL mL^-1^85% lactic acid, and 5 µg mL^-1^ hemin and liquid starter cultures were grown in the corresponding liquid formulation. All starter cultures were incubated under anaerobic conditions (5% H_2_, 5% CO_2_, and 90% N_2_).

### Screening for individual species or species interactions affecting Pg growth using factorial design of experiments (DOE) methodology

A previously developed 6-species polymicrobial community model^14^ in which Pg, inoculated at low density, is capable of growth in batch culture in mucin-serum medium with other 5 commensals was used as a basis to evaluate inter-species interactions that could influence Pg growth. For this purpose, we utilized the Design of Experiments (DOE) methodology. A half-fraction factorial experiment was designed using Design-Expert software (Stat Ease, Inc) allowing systematic testing of the effect of the five oral commensals (Ao, Vp, Ss, Fn and Rd) on Pg growth rate as outcome. The experiment included 16 distinct species combinations, representing half of all possible combinations and a Pg alone control. Communities were grown in batch culture anaerobically in mucin-serum medium, which contained 2.5 mg mL^−1^ hog gastric mucin (Sigma), 2.5 mg mL^−1^ KCl, 2.0 mg mL^−1^ proteose peptone, 1.0 mg mL^−1^ yeast extract, 1.0 mg mL^−1^ trypticase peptone, 1.0 μg mL^−1^ cysteine ·HCl, 5 μg mL^−1^ hemin and 10% (v/v) heat-inactivated human AB serum. All species, including Pg, were inoculated at a standardized starting density of 10^5^ cells mL^−1^. Cultures were sampled daily for 4 days. The biomass of Pg within the consortia was quantified via qPCR (see below) and Pg growth rate during exponential phase was calculated. Growth of other species in the consortium was monitored via plating on differential agar media. Specifically, Vp and Fn were plated on BHI-cysteine blood agar supplemented with 2.5 μg mL^−1^ vancomycin under anaerobic condition. Ao and Rd were plated on the Actinomyces selective agar containing 30 mg mL^−1^ trypticase soy broth, 5 mg mL^−1^ glucose, 13 μg mL^−1^ cadmium sulfate, 80 μg mL^−1^ sodium fluoride, 1.2 μg mL^−1^ neutral acriflavine and 0.25 μg mL^−1^ basic fuschin. Ao colonies were counted after anaerobic incubation, while Rd was quantified aerobically by subtracting morphologically distinct Ao colonies (raised, orange pigmentation) from total viable aerobic counts. Lastly, Ss was selectively enumerated on solid mitis salivarius agar. This screening experiment was performed in two independent days with technical duplicates included for each condition. Data were analyzed using Design-Expert software software (Stat-Ease, Inc), calculating a linear regression model to evaluate the main effects of individual species treated as a binary factor (present or absent) and their interaction effects on Pg growth rate. Multiple comparisons were corrected using the Bonferroni method, with statistical significance set at an adjusted p < 0.05.

### Co-culture of Pg with Ao, Ao mutant strains, Ao spent media and/or Vp

All experiments involving liquid co-cultures of Pg, Ao, and Vp strains were performed in mucin-serum medium. Microorganisms were either inoculated at low-density (10^5^ cells mL^−1^), at which Pg cannot grow alone but depends on Vp, or at high cell-density (10^7^ cells mL^−1^), at which Pg can grow independently^14^. Cultures were incubated anaerobically at 37°C. Culture samples were collected daily or every other day inside the anaerobic chamber. The pH of the collected culture samples was immediately measured in supernatants, collected after a brief centrifugation (8,000 × g for 4 min), via an Accumet AB15 pH meter. For quantification of Pg biomass, DNA was extracted from pellets previously stored at −80°C using the Qiagen Blood & Tissue Kit. qPCR was carried out using validated forward and reverse primers, along with SYBR Green detection, as previously described^47^. To collect cell-free culture supernatants, Ao strains were grown under aerobic or anaerobic conditions to the time point where each culture reached late exponential phase. Cultures were pelleted by centrifugation (8,000 × g, 10 min, 4°C), and supernatants were collected and sterilized via filtration through 0.22 μm polyethersulfone (PES) membrane syringe filters twice. Then filtered spent media were stored at −80°C until further use. Spent media supplementation assays were prepared by mixing cell-free supernatants with fresh mucin-serum medium at a 1:1 (v/v) ratio. Pg was inoculated at a final concentration of 10^7^ cells mL^−1^ and incubated anaerobically at 37°C. For pH-buffered conditions, the culture medium ingredients were added to previously prepared 10 mM potassium phosphate buffer (pH 7.3), instead of water, and the medium was autoclaved and subsequently supplemented with hemin, menadione and inactivated human AB serum (10% v/v). All quantitative analyses were performed across three independent biological experiments.

### Competitive spot assays

Pg and Ao starter cultures were grown under anaerobic conditions until they reached an optical density (600 nm) of 0.8, then normalized with fresh medium to a concentration of 10^7^ cell mL^−1^. 15 µL of a normalized Ao suspension were spotted on BHI or blood agar plates supplemented with hemin and menadione. 15 µL of a normalized Pg culture were spotted adjacent to Ao either simultaneously or after 24 hours of Ao incubation aerobically or anaerobically. The spots of Ao and Pg were placed close to each other but without their boundaries touching. The spotted agar plates were then incubated anaerobically for 72 hours at 37°C to allow for Pg growth. The area of Pg inhibition was calculated via ImageJ software. Two technical replicate spots were included in each experiment, and three independent experiments were performed.

### Phenol-red agar acidification analysis

To quantify the acidogenic activity of Ao on agar, the surface area of pH-induced color change was analyzed using ImageJ software via a standardized thresholding protocol. Phenol red agar was prepared by supplementing BHI agar with 18 mg L^−1^ phenol red (Sigma-Aldrich). To ensure optical consistency and uniform diffusion gradients, a precise volume of 25 mL was dispensed into each 100 mm petri dish. Following standardized concentration of Ao (10^7^ cells mL^−1^) spotting and a 24-hour incubation under aerobic or anaerobic conditions, digital images were captured under fixed illumination and focal distance. Spatial calibration was performed for each frame using the known internal diameter of the petri dish as a reference scale (pixels/mm). Acidified regions were isolated using the color threshold tool in the hue, saturation, and brightness (HSB) color space. Threshold parameters were defined by sampling a pre-calibrated pH standard series (pH 4.0–8.0), specifically setting the hue range to encompass the yellow chromophore characteristic of pH < 6.0 while excluding the magenta/red tones of the alkaline standards (pH 7.0). After applying these parameters to the experimental images, the selected regions were converted into binary masks, and the analyze particles function was employed to calculate the total integrated area (mm²) of acidification. Two technical replicate spots were included in each experiment and three independent experiments were performed.

### Quantification of organic acids by colorimetric assays

Extracellular concentrations of lactate, succinate, fumarate, and pyruvate were quantified using the L-lactate assay kit (Cell BioLabs, cat. no. MET-5012), Succinate assay kit (Abcam, cat. no. ab204718), EnzyChrom Fumarate assay kit (BioAssay Systems, cat. no. EFUM-100), and Pyruvate assay kit (Abcam, cat. no. ab65342). Ao wild-type and mutant strains were cultured in mucin-serum medium for 24 hours, and 1,100 μL culture aliquots were harvested. 100 μL of each aliquot was reserved for colony-forming unit (CFU) count, while the remaining volume was centrifuged to collect the supernatant (8000 × g, 10 min, 4°C). The resulting supernatants were sterilized by passing twice through 0.22 μm syringe filters and serially diluted in assay buffers. Diluted samples and standard curve serial dilutions were loaded in technical triplicates into flat-bottom 96-well microplates. Following the addition of enzyme reaction mixtures and incubation (either for 30 min at room temperature or at 37 °C, according to the manufacturer’s instructions), colorimetric absorbance was measured using a Synergy HT microplate reader (BioTek). Metabolite concentrations were calculated from corresponding standard curves and normalized to Ao viable cell counts (CFU mL^-1^). All quantitative analyses were performed across three independent biological experiments.

### Quantification of organic acids in spent media via targeted liquid chromatography-mass spectrometry (LC-MS)

Cell free culture supernatant collected as described above was also used for quantification of five organic acids via targeted LC-MS. Reference grade standards of sodium acetate, succinic acid, valeric acid, butyric acid, propionic acid, fumaric acid, sodium L-lactate, 1-^13^C-sodium acetate, 3-^13^C-lactic acid, 1-^13^C-butyric acid and 2,3-^13^C_2_-succinic acid were purchased from SigmaAldrich. All stock standard solutions were prepared at a concentration of 1 mg mL^-1^ in water. Chromatographic separation was carried out on an UltiMate 3000 HPLC coupled to a Q Exactive Focus Orbitrap mass spectrometer (Thermo Fisher Scientific) using a ThermoScientific Accucore^TM^-150-Amide-HILIC column (2.6 μm, 2.1mm x 150mm) maintained at 30°C. Calibration standards were prepared in acetonitrile at concentrations of 50, 25, 12.5, 6, 3, 1.5, and 0.78□μg mL^-1^ through serial dilution. Each standard was spiked with internal standards to achieve a final concentration of 19□μg mL^-1^. Spent media preparation involved double filtration using 0.2□μm syringe filters, followed by dilution of 100□μL of the filtrate to 1□mL in acetonitrile and a subsequent filtration through a 0.45□μm filter. Additional dilutions (either 2× or 10×) in acetonitrile were performed as needed during sample processing. Analytes were resolved in negative electrospray ionization mode using an isocratic mobile phase of 22% acetonitrile and 78% 20 mM ammonium formate buffer (pH 8.5) at a flow rate of 0.25 mL min^-1^ over an 8-min run with a 12 µL injection volume. Analyte peak identities were validated against individual pure reference standards to eliminate ambiguous assignments arising from structurally similar isomeric or isobaric fragments in extracted ion chromatograms (EICs). Standards and samples were injected in quadruplicate, interspersed with solvent blanks to verify absence of carryover. Data processing, unweighted linear regression, and analyte quantification were performed using TraceFinder software (v.4.1, Thermo Fisher Scientific) across three independent biological replicates.

### Genomic loci identification and sequence analysis

Annotated complete genome sequences of *Actinomyces oris* MG1 were retrieved from the NCBI GenBank database (accession no. CP194891.1). Genomic neighborhood maps spanning candidate metabolic loci with genes of interest were extracted, annotated, and visualized using SnapGene software (v8.2.2, Dotmatics). Multiple sequence alignments of target nucleotide and amino acid sequences were performed using MUSCLE (default parameters) implemented in MEGA12. Pairwise sequence divergence was determined using the uncorrected p-distance model, with missing data, ambiguous sites, and alignment gaps resolved via the pairwise deletion treatment option.

### Construction of Ao mutant strains and growth evaluation

In the genome of Ao, we identified two L-lactate dehydrogenase genes, a putative NAD+-dependent enzyme [*ldh1*, ACSAM2_10450, EC 1.1.1.27] and a quinone-dependent enzyme [*ldh2*, ACSAM2_00080, EC 1.1.5.15] as well as two distinct succinate dehydrogenase/fumarate reductase heterotrimeric complexes (EC 1.3.5.1), each having a distinct catalytic flavoprotein subunit [*sdh*1, ACSAM2_04915] and [*sdh*2, ACSAM2_12050]. The in-frame deletion mutants Δ*ldh1*, Δ*ldh2,* Δ*sdh1*, Δ*sdh2* and the double mutants (Δ*ldh1/*Δ*ldh*2, Δ*sdh1/*Δ*sdh2*, Δ*ldh1*/Δ*sdh1* and Δ*ldh1/sdh2)* were generated using a previously published protocol.^48^ Briefly, 1 kb regions upstream and downstream of *ldh* or *sdh* genes were PCR-amplified and joined together using overlap extension PCR with appropriate primers. The 2 kb fragments were cloned into the deletion vector pCWU2. Competent cells of an Ao Δ*galK* strain were transformed with each deletion plasmid using electroporation to obtain kanamycin-resistant integrant strains^49^. The strains were grown overnight without kanamycin that facilitated the second homologous recombination event, thereby leading to plasmid excision. The cells were then diluted and grown on agar plates containing 2-deoxy-D-galactose to screen for mutant colonies. The mutants were verified by colony PCR and DNA sequencing. All Ao cultivation during the mutagenesis process was performed under aerobic conditions. The fitness of Ao mutant strains under different redox conditions was evaluated in mucin-serum and BHI media. For aerobic growth, cultures were incubated at 37°C with shaking at 200 rpm. Microaerophilic cultures were grown statically at 37°C in a CO_2_incubator, and anaerobic cultures were maintained at 37°C in an anaerobic chamber (5% H_2_, 5% CO_2_, 90% N_2_). Bacterial growth was monitored at indicated time points by collecting 10 μL culture aliquots, preparing serial dilutions, plating onto BHI agar, and enumerated via colony-forming units (CFU mL^-1^). Culture pH was monitored by harvesting 200 L aliquots at each time point, centrifuging at 8000 × g for 4 min, and measuring the supernatant pH using an Accumet AB15 pH meter. For spent media supplementation experiments, cultures of wild-type and mutant strains grown under aerobic or anaerobic conditions were harvested during late logarithmic phase. Cell-free supernatants were collected by centrifugation (8000 × g, 10 min), filter-sterilized (0.22μm PES membrane), and mixed 1:1 (v/v) with fresh mucin-serum medium followed by Pg inoculation.

### Human subgingival microbiome analysis

To assess whether the interactions among Ao, Pg and Vp observed in vitro were reflected in human subgingival communities, we analyzed previously generated subgingival 16S rRNA gene sequencing data from 1,355 adults (588 women and 767 men) aged 35–72 years. This cohort included 534 survivors of incident non-fatal myocardial infarction and 821 community controls^31^. Recruitment was independent of periodontal status, and participants spanned the spectrum from no periodontitis to severe periodontitis. Participants had at least six natural teeth and individuals reporting systemic antibiotic use within one month before examination were excluded. The study was approved by the Institutional Review Boards of the University at Buffalo and participating hospitals in Erie and Niagara counties, New York (IRB #030-488953), and all participants provided written informed consent. Periodontal status was classified using the Centers for Disease Control and Prevention/American Academy of Periodontology (CDC/AAP) surveillance definitions ^32^. Severe periodontitis required ≥2 interproximal sites on different teeth with CAL ≥6 mm and ≥1 interproximal site with PD ≥5 mm. Moderate periodontitis required ≥2 interproximal sites on different teeth with CAL ≥4 mm, or ≥2 interproximal sites on different teeth with PD ≥5 mm. Mild periodontitis required ≥2 interproximal sites with CAL ≥3 mm together with either ≥2 interproximal sites on different teeth with PD ≥4 mm or ≥1 interproximal site with PD ≥5 mm. Participants were assigned to the most severe category for which they met the criteria; those meeting none were classified as having no periodontitis, labelled Healthy in the figures. Subgingival plaque was collected from the mesiobuccal sulcus or pocket of 12 preselected index teeth using paper points inserted for 10 s; prespecified substitute teeth were sampled when index teeth were absent. Maxillary and mandibular samples were stored in liquid nitrogen and combined to generate one pooled sample per participant. The microbiome was characterized via amplicon sequencing of the V3–V4 regions of the bacterial 16S rRNA gene. Reads were processed using a pipeline that included DADA2^50^. Amplicon sequence variants (ASV) were annotated against the Human Oral Microbiome Database (v15.22) ^51^ using BLAST ^52^, with species assignments accepted at ≥99% sequence similarity; unresolved sequences were further classified using the Ribosomal Database Project classifier ^53^. We used the resulting taxonomic relative-abundance profiles for the present analysis. Detailed laboratory and sequence-processing procedures are reported in Li et al.^31^. Sequence data are available in the NCBI Sequence Read Archive under accession PRJNA1430094.

### Identification of fermentation genes in Actinomyces genomes

To assess the genomic conservation of key organic acid fermentative genes across *Actinomyces* species, representative whole-genome sequences corresponding to each identified species and HMT strain in our dataset were retrieved directly through the HOMD platform. The presence or absence of genes within the lactate dehydrogenase and succinate dehydrogenase branches was then evaluated. Candidate loci for *ldh1*, *ldh2*, *sdh1* and *sdh2* were identified by cross-referencing curated NCBI GenBank annotations with standardized PROKKA annotations in the HOMD Genome Table viewer. To verify orthology, translated amino acid sequences of candidate genes from each species were retrieved and aligned against verified Ao reference sequences via multiple sequence alignment using MUSCLE in MEGA. Pairwise genetic distances ( -distance) were calculated to confirm sequence identity. Validated gene presences were compiled across all evaluated taxa into a categorical presence–absence matrix for visualization.

### Statistical analysis

All quantitative experiments were conducted across at least three independent biological replicates, each evaluated in duplicate or triplicate technical replicates. Differences among experimental groups were evaluated using one-way analysis of variance (ANOVA) followed by Fisher’s least significant difference (LSD) test, or using the Kruskal–Wallis test followed by Dunn’s test for nonparametric pairwise comparisons. Unpaired two-tailed Student’s t-tests were used for two-group comparisons where appropriate. We corrected for multiple comparisons by controlling the false discovery rate using the Benjamini–Hochberg method. All quantitative results are presented as mean standard deviation (SD), with error bars indicating SD. Statistical significance was defined as *P* < 0.05.

Microbiome association analyses were restricted to Pg-positive subgingival plaque samples (n = 555). Samples were divided into low-, medium-and high-Vp groups using tertiles of Vp relative abundance. The low and medium tertiles were combined into a low/medium-Vp group (n = 370), while the upper tertile constituted the high-Vp group (n = 185). Pg, Ao and genus-level *Actinomyces* relative abundances were expressed as percentages (0–100%). A pseudocount of 0.0001 was added to each value before log transformation for correlation and regression analyses. Pearson correlation coefficients were calculated separately within the two Vp groups, and unadjusted linear fits were plotted for visualization. Separate multiple linear regression models were fitted for Ao and genus-level *Actinomyces*, with transformed Pg abundance as the outcome. Each model included transformed Ao or genus *Actinomyces* abundance, Vp group, their interaction and CDC periodontal disease group as a categorical covariate. Vp group was coded as 0 for low/medium Vp and 1 for high Vp. The taxon abundance coefficient therefore represented its association with Pg in the low/medium-Vp reference group, while the interaction coefficient estimated the difference in slope between the two Vp groups. Regression coefficients (β), 95% confidence intervals and two-sided *P* values were reported.

## Acknowledgements

This work was supported by grants R21DE023967 and R21DE034093 (to P.I.D.) and R01DE017382 (to H.T.-T.) from the National Institute of Dental and Craniofacial Research, National Institutes of Health. Dam Soh was supported by grant K12 DE027827 and Lu Li by grant K99DE034829 from the National Institute of Dental and Craniofacial Research, National Institutes of Health.

## Data availability

The microbiome sequence data have been deposited in the NCBI Sequence Read Archive under BioProject accession PRJNA1430094. Associated demographic and clinical metadata are available in the linked BioSample records.

