## Supplementary Figures for "Higher-order metabolic interactions among oral commensals gate pathobiont expansion through succinate-driven pH modulation"

Supplementary Figure 1. (a) Growth of oral commensal species, *Veillonella parvula* (Vp), *Actinomyces oris* (Ao), *Streptococcus sanguinis* (Ss), *Fusobacterium nucleatum* (Fn), and *Rothia dentocariosa* (Rd), across all conditions in factorial experiment that included each microorganism. Biomass after days 1 and 2, quantified by CFU is shown. (b) Pareto chart of standardized effects on Pg specific growth rate ($h^{-1}$). Absolute standardized effects ($|t|$) are ranked for main effects and two-factor interactions. The grey and red horizontal lines indicate the uncorrected ($t = 2.18$, $\alpha = 0.05$) and Bonferroni-adjusted ($t = 3.61$) significance thresholds, respectively. Orange and blue bars denote positive and negative effects. (c) Viable cell counts of Vp grown in mucin–serum medium in co-culture with Pg or with Pg and Ao. Error bars in a and c indicate standard deviations.

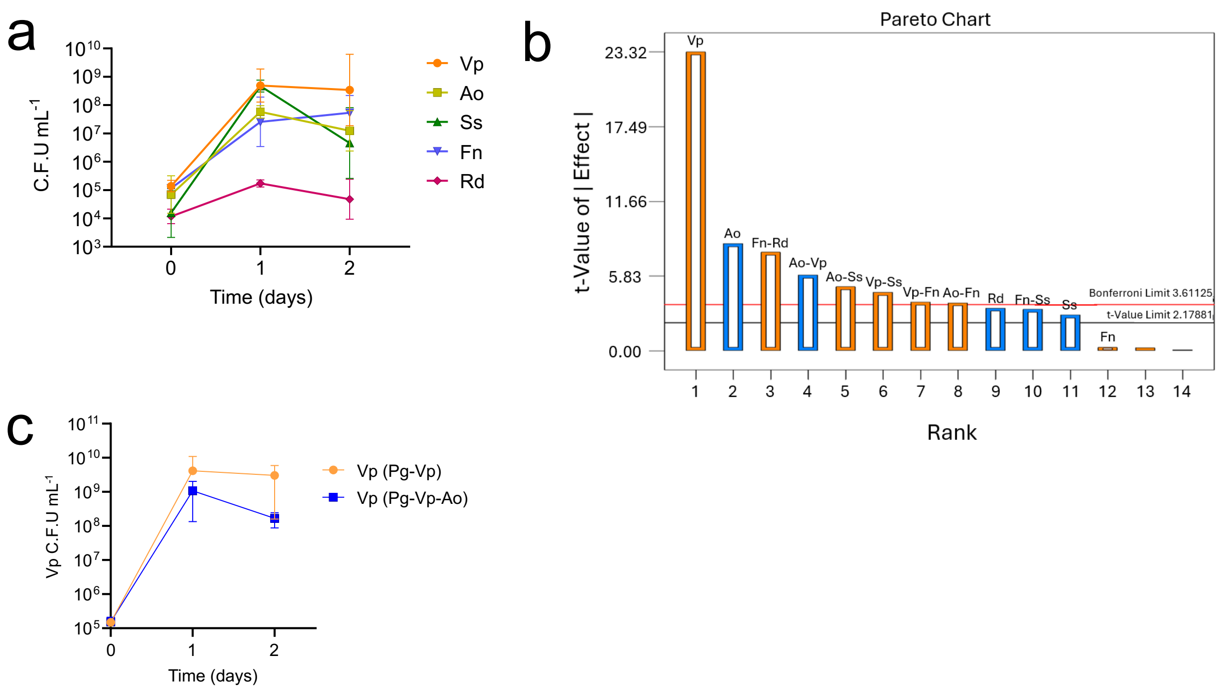

Supplementary Figure 2. Predicted oxygen-dependent rewiring of the Embden–Meyerhof–Parnas (EMP) pathway and organic acid branch points in Ao. Schematic illustrates predicted fermentative pathways for glucose utilization in Ao under aerobic versus anaerobic conditions.

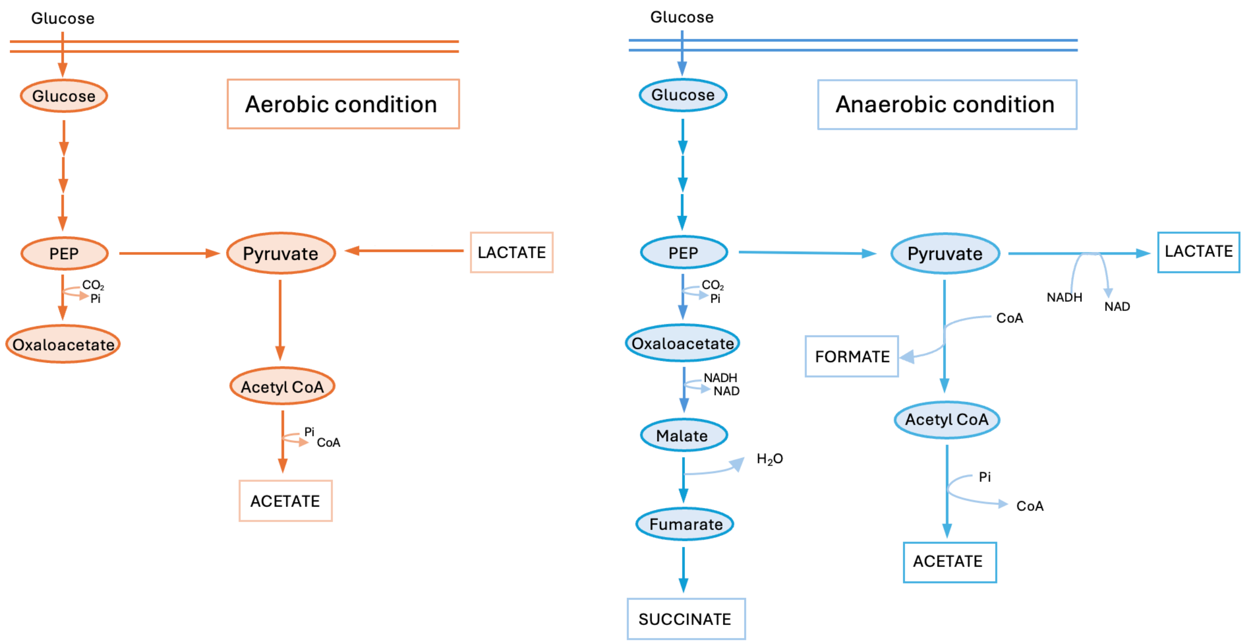

Supplementary Figure 3. Deletion of *ldh* genes alters Ao growth kinetics and medium acidification in rich medium. Growth kinetics (optical density, upper panels) and culture pH (lower panels) of Ao WT and lactate dehydrogenase (*ldh*) mutant strains cultured in brain heart infusion (BHI) broth under aerobic or anaerobic conditions over time.

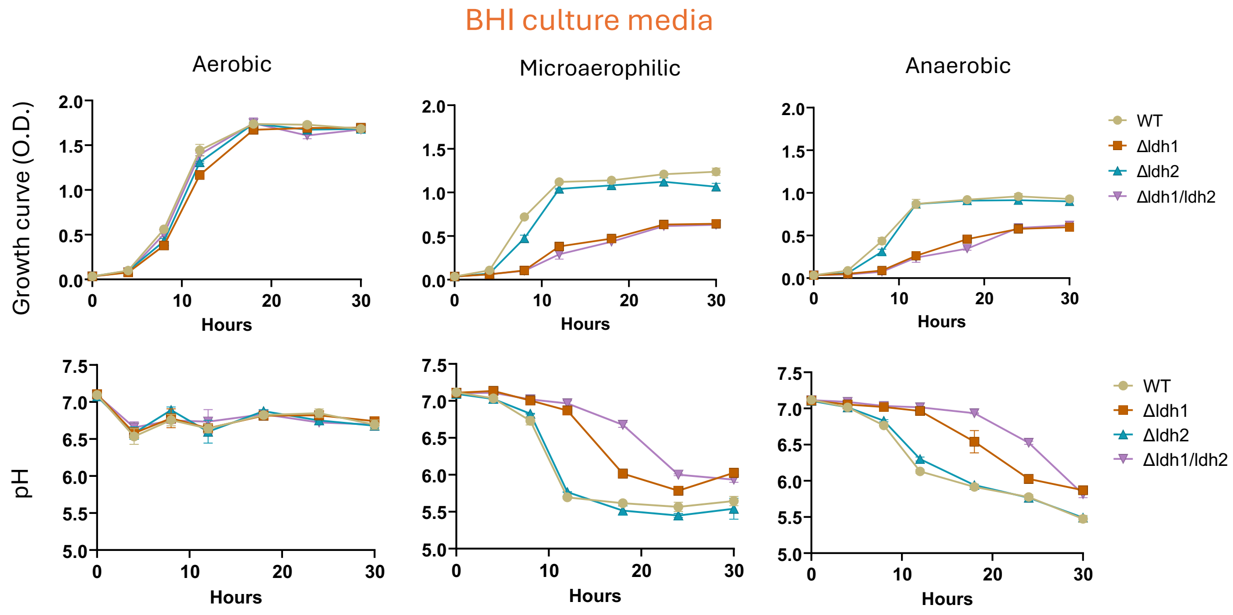

Supplementary Figure 4. Supplemental information for metabolic profiles of Ao *sdh* mutants and their inhibitory capacity against *Pg*. (a) Extracellular pyruvate accumulation in cell-free supernatants of Ao WT and *sdh* mutant strains cultured anaerobically in mucin–serum medium for 24 hours. (b) Extracellular succinate accumulation in cell-free supernatants of Ao WT and *sdh* mutant strains cultured aerobically in mucin–serum medium for 24 hours. Pyruvate (a) and succinate (b) were quantified by enzymatic colorimetric assays and normalized to total cell counts. (c) Pg growth inhibition in adjacent spot competition assays on BHI blood agar using Ao WT and *sdh* mutant strains pre-grown aerobically for 24 hours. (d) Growth of Pg inoculated in mucin–serum medium supplemented with cell-free spent medium from aerobic Ao WT or *sdh* mutant cultures. Pg monoculture and uninoculated medium provide growth and pH references, respectively. The left panel shows Pg abundance quantified by qPCR and the right panel displays corresponding culture pH. Error bars indicate standard deviations. Differences among strains in a–c were assessed using one-way ANOVA followed by Tukey's multiple comparisons test. * $P < 0.05$, ** $P < 0.01$, *** $P < 0.001$.

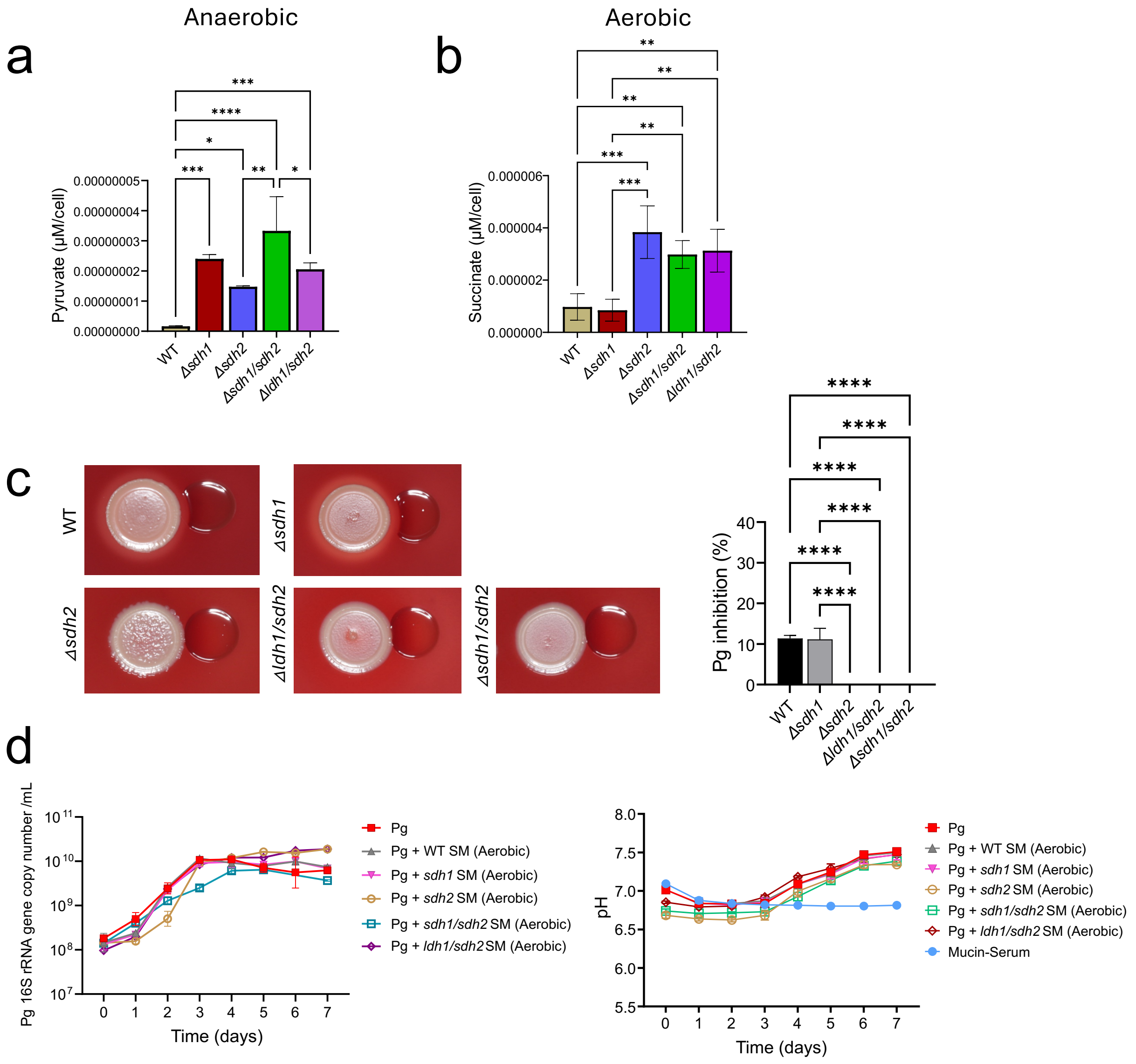

Supplementary Figure 5. Population-level prevalence, abundance, and conditional associations of Pg, Ao and Vp (a) Prevalence of Pg, Ao, and Vp in subgingival plaque samples from participants across periodontal disease severity groups (n = 1,355), determined by 16S rRNA gene sequencing. (b) Relative abundance of genus *Actinomyces* across the same subgingival plaque samples described in a. (c) Interaction plot showing Pg prevalence in Ao-negative and Ao-positive samples, stratified by Vp presence (pink square) or absence (blue circles). Pg prevalence was compared between Ao-negative and Ao-positive samples within each Vp stratum using Fisher's exact tests.

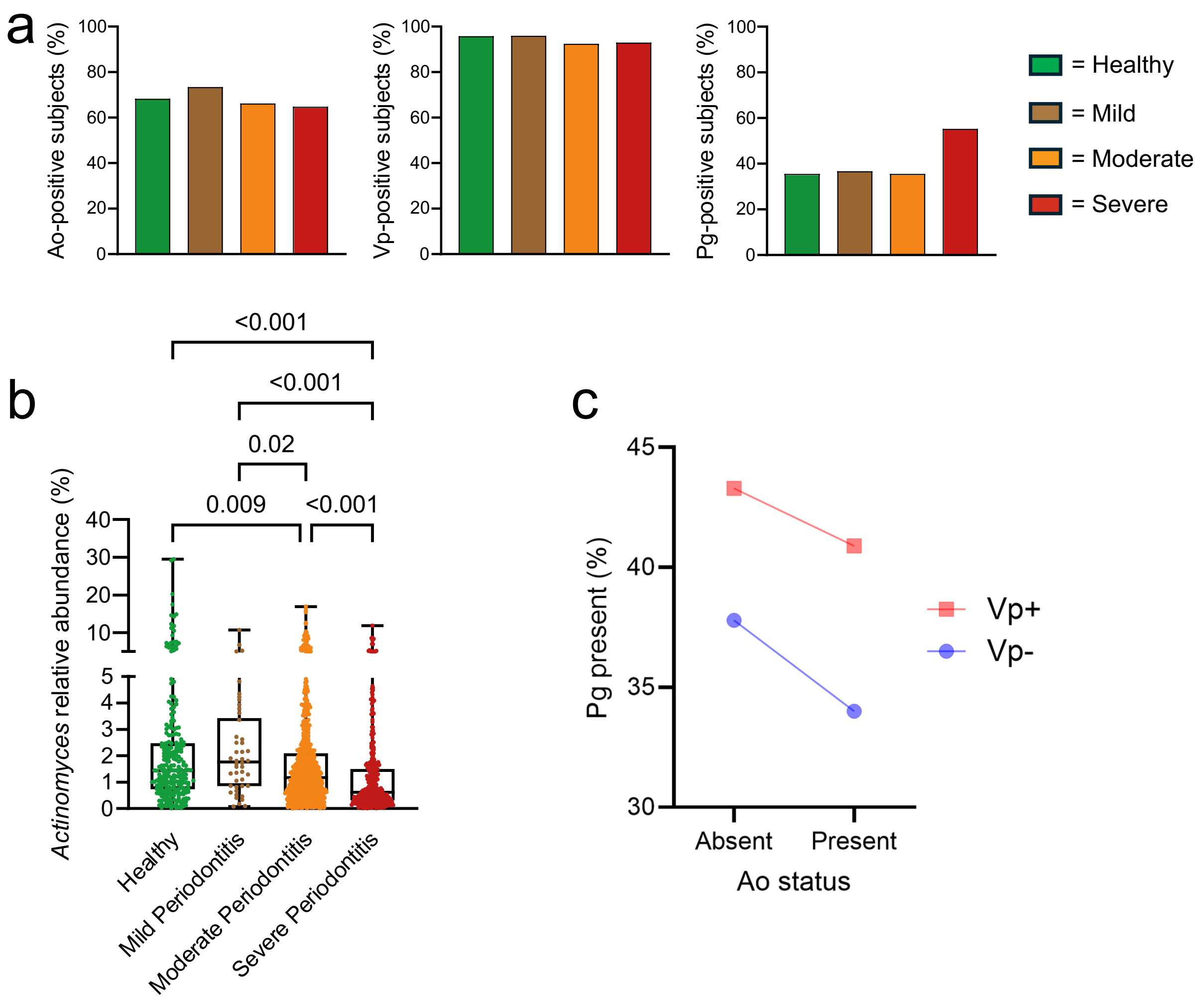
